# Gradual consolidation of skilled sequential movements in primary motor cortex of non-human primates

**DOI:** 10.1101/2024.10.06.616850

**Authors:** Machiko Ohbayashi, Nathalie Picard

**Author notes:** Corresponding author: Machiko Ohbayashi.

## Abstract

Many of our daily actions rely on skilled sequential movements. Sequence performance improves with practice and can reach expert levels with years of repeated practice, truly exemplifying the saying “Practice makes perfect”. Human and non-human primate studies have shown structural and functional changes in the primary motor cortex (M1) following extended practice, suggesting M1’s involvement in acquisition and retention of skilled sequential movements. However, it has been challenging to causally examine M1’s role in sequence learning, because inactivation or lesion of M1 impairs movement production itself. Here, we causally examined M1’s contribution to learning of sequential movements by locally inhibiting protein synthesis in M1. Our results show that protein synthesis inhibition in M1 disrupted memory-guided sequential movements at all stages of learning while sparing visually-guided reaching accuracy, though the effects decreased with continued practice. These findings suggest that neural representations of sequential movements are repeatedly consolidated in M1 through protein synthesis, with the rate of consolidation slowing as learning progresses.

**Teaser:** Repetitive practice strengthens neural representations of sequential movements in primary motor cortex of monkeys.

## Introduction

Many daily activities, such as playing a musical instrument and typing, rely on attaining a high level of proficiency in performing sequential movements. Sequential movements can be acquired and improved to an expert level through extensive practice, eventually becoming a stable motor skill. Once the structure of a sequence is learned during the early phase of learning, continued daily practice over an extended period leads to gradual improvement in speed and accuracy, key indicators of motor skill learning. This slow incremental learning is considered a fundamental aspect of motor skill development, capturing the idea that “practice makes perfect.”

Emerging evidence indicates that multiple cortical motor areas alter their activity during the learning of sequential movements (*1–8*). The primary motor cortex (M1), in particular, exhibits functional and structural changes associated with the learning of sequential movements, not only during the early stages of learning but also after extensive practice. In humans, short-term training induces measurable changes in fMRI activity within M1 (*1–7, 9, 10*). Similarly, studies in monkeys demonstrate that neural activity in M1 is modulated by sequence components during the early stage of learning (*11, 12*).

With extensive practice, additional changes in M1 have been reported in both human and non-human primate. In humans, years of extensive practice are associated with structural modifications and altered functional activation of M1 (*7, 8, 13–17*). For instance, professional musicians who have spent years practicing complex sequential movements exhibit reduced or more focused M1 activation during the performance of sequential tasks compared with non-musicians or amateurs (*13–17*). This reduced activation is considered as evidence of increased efficiency of neural circuits, or the need for a smaller number of active neurons to perform a highly trained set of sequential movements. Furthermore, the volume of M1 has been reported to be larger in professional musicians compared to amateurs or non-musicians (*18–24*). These structural changes have been proposed to arise from synaptic processes, including intracortical remodeling of dendritic spines and axonal terminals, glial hypertrophy, and synaptogenesis as a result of extensive practice (*21, 22, 25*). Similarly, plasticity of the white matter structure of M1 has been correlated with the amount of time devoted to practice, suggesting that myelination may be increased by neural activity in fiber tracts during training (*26, 27*).

In non-human primates, after years of extensive training, M1 neurons were differentially active during the performance of visually guided and internally generated sequential reaching (*28, 29*). Furthermore, the uptake of 2-deoxyglucose (2DG), a measure associated with presynaptic activity, was lower in M1 of monkeys that performed highly practiced, internally generated sequences of movements compared with M1 of monkeys that performed visually guided reaching (*29*). Changes in M1 observed at the levels of neuronal activity, functional activation, and gray matter structure in humans and non-human primates may all reflect plasticity underlying the learning of sequential movements through extended, repetitive practice.

Together, these diverse findings converge to the concept that M1 contributes to the learning and maintenance of motor skills, such as sequential movements through extensive practice. While these studies demonstrate that activity patterns in M1 change both during the early phase of learning and after extensive practice, they do not conclusively indicate whether M1 is critical for sequence learning nor when and how plasticity occurs within M1 during learning. The change after extensive training could be the outcome of plasticity during the early phase of the training. Thus, it remains unclear whether M1 undergoes changes throughout all phases of learning or only during specific stages, and whether M1 plays a critical role in learning of sequential movements. Moreover, the mechanisms by which M1 contributes to the slow, gradual improvements in sequence performance are not yet fully understood.

A traditional approach to these questions would be to assess the effect of lesion or inactivation of M1 on learning performance. However, M1 poses a special challenge to causally examine when and how plastic changes occur during learning, because M1 is critical for implementing motor output. A lesion or inactivation of M1 will abolish the motor commands to the spinal cord that generate muscle activity, so that it would be impossible to evaluate M1’s involvement in learning. Therefore, we targeted information storage in M1 instead of motor output. We injected an inhibitor for protein synthesis in M1 to interfere with information storage and, consequently, learning.

Inhibitors for protein synthesis have been widely used to dissect and analyze memory systems in rodents, such as fear conditioning. Local injection of an inhibitor of protein synthesis (e.g., anisomycin) into amygdala, hippocampus or motor cortex of rodents disrupted the learning, maintenance or reconsolidation of memory (*30–39*). These studies indicated that protein synthesis inhibition can interfere with formation and storage of memory. Building on this approach, we previously injected a protein synthesis inhibitor into M1 of monkeys after ∼100 days of training and found that it successfully disrupted performance of well-trained sequential movements without affecting the performance of visually guided reaching (*40*). Here, we applied this approach to examine when and how M1 contributes to learning of sequential movements during repetitive practice over extensive time in non-human primates.

## Results

### Learning of sequential reaching task

We trained three monkeys (*Cebus apella*) to perform Random and Repeating tasks in alternating blocks (Fig. 1A–E; see Materials and Methods for additional details). The monkey sat in front of a touch-sensitive monitor with the left arm restrained. Five square targets were outlined on the screen (Fig. 1A, B). In both tasks, the monkey made a series of continuous reaches from one target to the next with the right arm, without an inter-trial interval. At the start of each trial, one target was filled with yellow which served as the visual instruction cue. To make a correct response, the monkey was required to touch the instructed target within 800 ms. The trial ended when the monkey contacted the screen or when 800 ms elapsed. Immediately after screen contact, the yellow cue disappeared and the next trial began. The two tasks used the same target array and movements but differed in the timing and order of instruction cues.

**Fig. 1.**
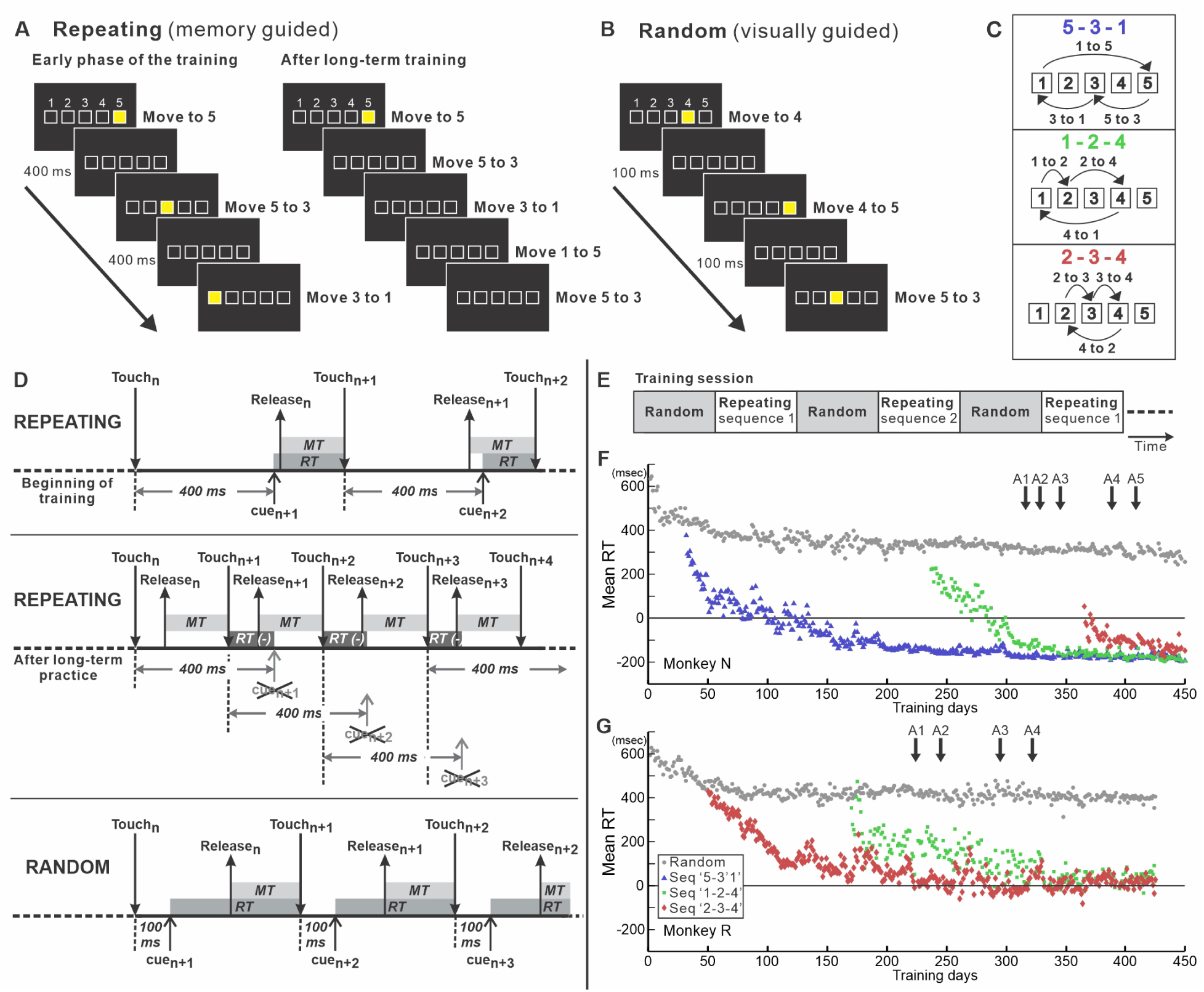
Sequential Reaching Task. **(A)** Repeating task. Visual cues are presented according to a predetermined sequence. During early learning, the monkey follows the visual cues and touches the targets in the instructed order (left). As learning progresses, the monkey begins to touch the next target in the sequence before the cue is presented. After extensive practice, the monkey memorizes sequences and performs them without relying on visual cues (right). **(B)** Random task. A monkey reaches a visual cue presented on a monitor 100 ms after touching the previous target. Cues are presented in a pseudorandom order, and the monkey reaches from one target to the next. **(C)** Three-element sequences in the Repeating task. **(D)** Task timing. RT, response time; MT, movement time. **Top:** Repeating task during early training. Visual cues are presented 400 ms after the monkey touches a target, following a predetermined sequence. **Middle:** Repeating task after extensive practice. The monkey predicts and touches the next target before the scheduled cue onset; the cue is not presented and the RT can be negative. **Bottom:** Random task. A visual cue is presented 100 ms after the monkey touches the monitor. The monkey is required to touch the target within 800 ms. **(E)** During training sessions, the monkey performed the Random and Repeating tasks in alternating blocks of 200–500 trials. **(F, G)** Response times (RTs) during Random (gray circles) and Repeating tasks for Monkey N (F) and Monkey R (**G**). Repeating sequences: ‘5-3-1’ (blue triangles), ‘1-2-4’ (green squares), and ‘2-3-4’ (red diamonds). Black arrows indicate anisomycin injections. Repeating sequences were introduced after ∼50 days of Random task training. Introduction of each sequence was staggered by 100–200 days. This training design allowed comparison of the injection effects on sequences with different training histories within the same injection session.

In the Repeating task, target locations followed a repeating three-element sequence (e.g., 5-3-1-5-3-1-…) (Fig. 1A, C). The instruction cue for the next target was scheduled to appear 400 ms after contact with the preceding target (Fig. 1D). If the monkey reached the correct next target during this 400-ms delay, the cue was omitted and the task advanced immediately to the following element in the sequence (Fig. 1A, D). With practice, the monkeys learned the sequence and increasingly initiated correct reaches before the scheduled cue appeared. These predictive responses therefore reflected memory-guided performance of the learned sequence. The monkeys could execute the sequence continuously without pause or cue.

In the Random task, which served as a control for motor performance, instruction cues were presented in pseudorandom order 100 ms after contact with the preceding target (Fig. 1B, D). Predictive responses were not possible and the visual cue guided reaching on each trial. Thus, the Random and Repeating tasks required similar reaching movements but differed in the mode of target selection, i.e. guided by memory for a learned sequence or by the current visual instruction.

During each daily session, the monkeys performed the Random and Repeating tasks in alternating blocks of 200–500 trials (Fig. 1E). After approximately 50 days of training, predictive responses occurred on more than 80% of Repeating task trials (Fig. 1F, G). Each monkey was trained on two or three Repeating sequences, introduced one at a time on a staggered schedule. A new sequence was introduced only after the preceding sequence had been practiced for more than 50 days (Fig. 1F, G; see Materials and Methods).

As training progressed, response times (RTs; measured as the time interval between two successive touches minus the imposed delay) decreased independently for each sequence (Fig. 1F, G), allowing us to examine the effects of pharmacological manipulations at different stages of learning within the same animal. For example, on day 408 in Fig. 1F, the three sequences had been practiced for 377, 171, and 42 days, respectively. In contrast, RTs in the Random task (gray dots in Fig. 1F, G) reached a plateau after approximately 20 days of training. This staggered training design therefore enabled us to compare the effects of pharmacological manipulations on sequences with different training durations and learning stages within a single injection session, as well as across multiple injection sessions over time.

### Protein synthesis inhibition disrupted memory-guided sequence performance while preserving visually guided reaching accuracy

When the monkey was able to perform the Random task and more than one Repeating sequence, we injected the protein synthesis inhibitor, anisomycin (100 mg/ml), into sites in M1 where intracortical microstimulation evoked shoulder or elbow movements (Fig. 2; see Materials and Methods). For each session, we compared task performance before and after injection, analyzing each component movement separately in both the Random and Repeating tasks (see Materials and Methods).

**Fig. 2.**
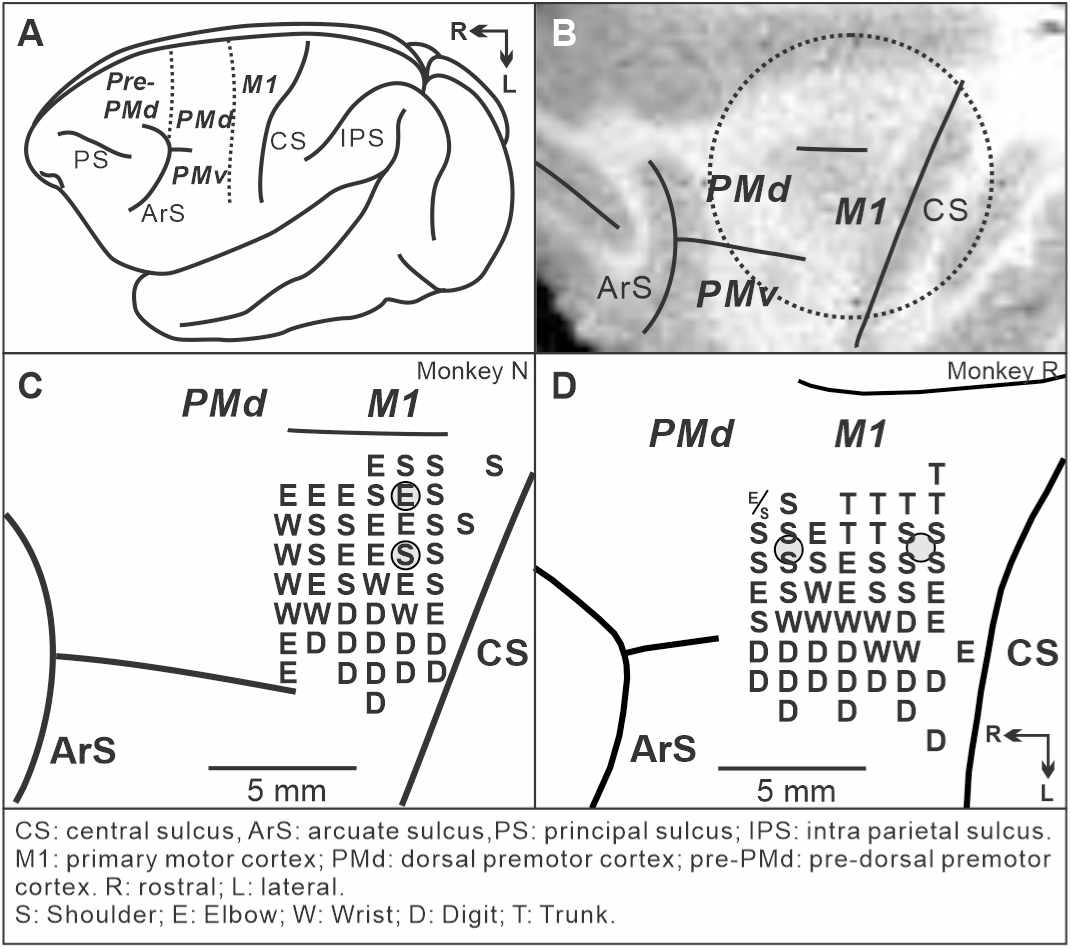
M1 location and intracortical microstimulation maps. **(A)** Schematic lateral view of the *Cebus* brain. Dashed lines indicate the approximate borders between M1 and PMd and between pre-PMd and PMd. PS, principal sulcus; ArS, arcuate sulcus; CS, central sulcus; IPS, intraparietal sulcus; pre-PMd, pre-dorsal premotor cortex; PMd, dorsal premotor cortex; M1, primary motor cortex. R, rostral; L, lateral. Adapted from Ohbayashi (2020) under CC BY 4.0. **(B)** MRI image after chamber implantation for monkey N. The dotted circle indicates the chamber outline. **(C, D)** Intracortical stimulation maps of monkey N **(C)** and monkey R **(D)**. Letters indicate the movements evoked by intracortical microstimulation at each M1 site. S, Shoulder; E, Elbow; W, Wrist; D, Digit; T, Trunk. Gray circles indicate representative anisomycin injection sites.

A representative session from Monkey N (session A5) illustrates the behavioral effects of anisomycin injection into M1 (Fig. 3; Supplementary Movies S1–S3). In this session, the monkey performed three sequences with different training durations: sequence 2-3-4 after 42 days of training (short-term; Fig. 3A), sequence 1-2-4 after 171 days (long-term; Fig. 3B), and sequence 5-3-1 after 377 days (extended; Fig. 3C). The anisomycin effect was particularly prominent for certain component movements within each learned sequence (Fig. 3D, F, H, J, L, N, P, R, T; Figs. S1 and S2). For example, the post-minus-pre changes in error rate were -19.25%, 5.78%, and 37.05% for movements ‘4-2’, ‘2-3’, and ‘3-4’ of sequence ‘2-3-4’; 34.16%, 6.89%, and 9.68% for movements ‘4-1’, ‘1-2’, and ‘2-4’ of sequence ‘1-2-4’; and 0.94%, 14.11%, and 30.48% for movements ‘3-1’, ‘5-3’, and ‘1-5’ of sequence ‘5-3-1’. Such movement-selective effects are consistent with the known movement-related organization and tuning properties of M1 neurons (*28, 41–43*). Despite this movement-to-movement variability, anisomycin produced a clear impairment when performance was evaluated at the level of the complete three-movement sequence (Fig. S3).

**Fig. 3.**
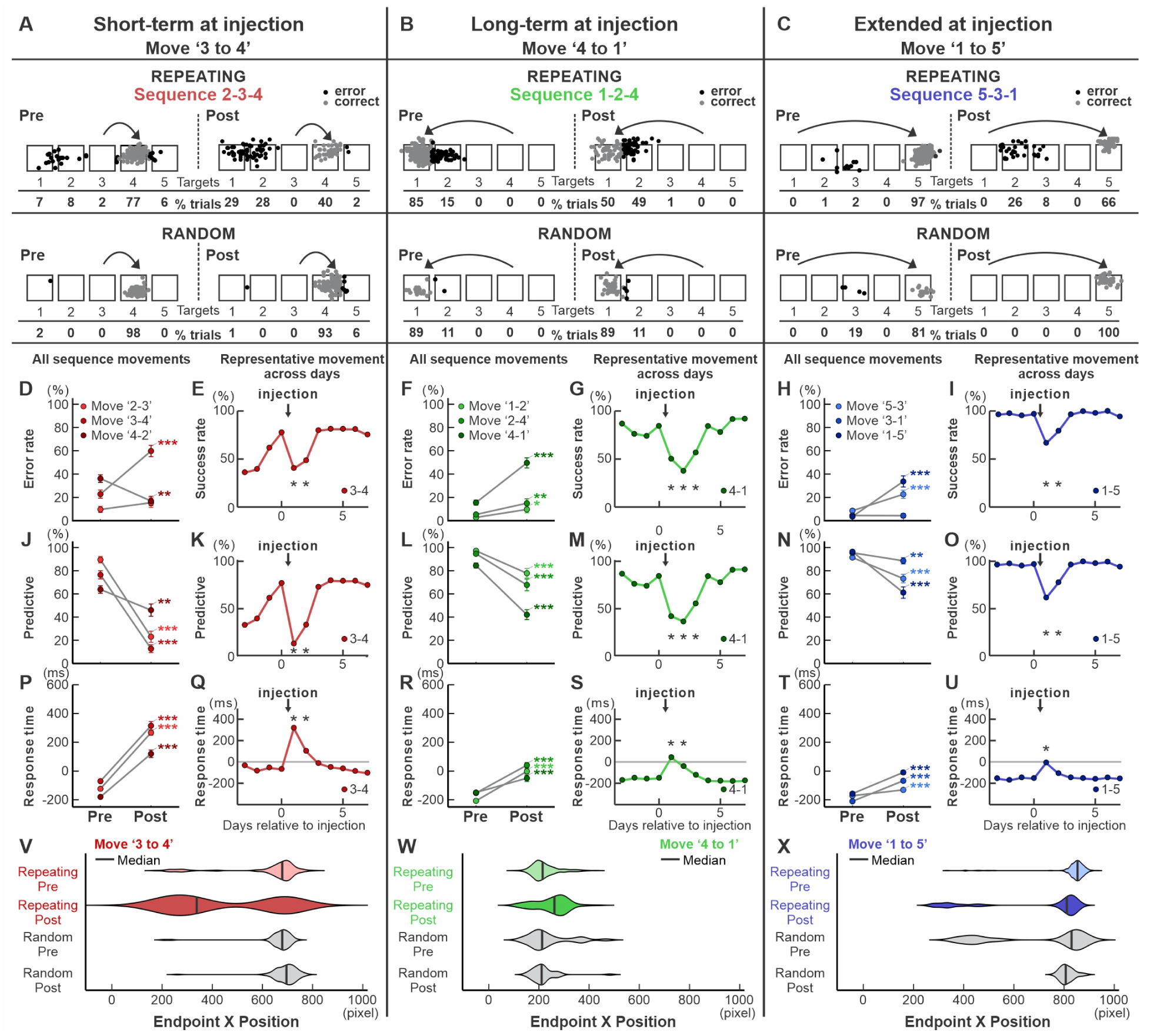
Effects of anisomycin injection on Repeating-and Random-task performance in a representative session (Monkey N, session A5). **(A, D, E, J, K, P, Q, V)** Short-term-trained sequence ‘2-3-4’ (42 days). **A**, Reach endpoints for the representative movement ‘3-4’ before and after anisomycin injection in the Repeating and Random tasks. Gray and black dots indicate correct and error responses, respectively; arrows indicate the instructed movement; percentages of responses to each target are shown below. **(D, J, P)** Pre- and post-injection error rate, predictive-response rate, and response time (RT), respectively, for all three component movements. Pre–post differences were tested separately for each movement using Fisher’s exact test for error and predictive rates and Welch’s two-sample *t*-test for RT. *P* values were Holm-adjusted across the three component movements separately for each metric. **(E, K, Q)** Daily time course of success rate, predictive-response rate, and RT for movement ‘3-4’. The arrow indicates the injection; each post-injection day was compared with the pre-injection baseline using χ² tests for success and predictive rates and two-sample *t*-tests for RT. **(V)** Endpoint-position distributions for movement ‘3-4’ in the Repeating and Random tasks; vertical lines indicate medians. **(B, F, G, L, M, R, S, W)** Corresponding analyses for the long-term-trained sequence ‘1-2-4’ (171 days), with movement ‘4-1’ shown in the representative panels. **(C, H, I, N, O, T, U, X)** Corresponding analyses for the extended-trained sequence ‘5-3-1’ (377 days), with movement ‘1-5’ shown in the representative panels. Asterisks in **D, F, H, J, L, N, P, R,** and **T** indicate Holm-adjusted significance (\**P* < 0.05, \*\**P* < 0.01, \*\*\**P* < 0.001). In the daily time-course panels, asterisks indicate significant differences from baseline (*P* < 0.05).

The pattern of errors indicated a disruption of learned sequence performance rather than a generalized inability to execute reaches. In this representative session, error rates increased in all three Repeating sequences after anisomycin injection (Fig. 3D, F, H; Fig. S1A-C). In contrast, percent-correct performance during visually guided movements in the Random task was not significantly impaired (Fig. 3A-C; Fig. S1D). Thus, the monkey retained the ability to select and execute visually instructed reaches despite the pronounced disruption of memory-guided sequence performance.

We further analyzed reach endpoints during the Repeating task to characterize the errors induced by anisomycin injection (Fig. S4). We first classified errors according to movement direction and amplitude (Fig. S4A–C). Direction Errors occurred when the monkey reached in a direction opposite to the correct target, whereas Amplitude Errors occurred when the monkey reached in the correct direction but failed to contact the intended target. For example, after anisomycin injection, the monkey moved in the wrong direction during movement ‘3-4’ of sequence ‘2-3-4’ in 57% of trials, whereas during movement ‘4-1’ of sequence ‘1-2-4’, the monkey moved in the correct direction but undershot the target in 50% of trials (Fig. 3A, B; Fig. S4A, B). Direction Errors increased significantly in sequence ‘2-3-4’, whereas Amplitude Errors increased significantly in sequences ‘1-2-4’ and ‘5-3-1’ (*P* < 0.001; Fig. S4A–C).

We also classified errors according to their relationship to the component movements of the sequence or their order (Fig. S4D–F). Out-of-Sequence errors occurred when the monkey reached to a target that was not part of the current sequence, whereas Off-Phase errors occurred when the monkey selected a sequence target at the wrong phase. Out-of-Sequence errors increased significantly in sequences ‘2-3-4’ and ‘5-3-1’, whereas Off-Phase errors increased significantly in sequences ‘1-2-4’ and ‘5-3-1’ (*P* < 0.001; Fig. S4D–F). Thus, anisomycin produced multiple types of errors in selecting and executing movements within the learned sequence rather than a single stereotyped deficit.

Anisomycin also impaired the predictive component of sequence performance. Before injection, the monkey frequently initiated the next correct movement before presentation of the visual instruction cue (>70% of Repeating-task trials; Fig. 3J, L, N). After injection, predictive responses decreased and RT increased (Fig. 3; *χ² tests* for predictive responses and *t-tests* for RT, *P* < 0.01). On some trials, the monkey paused during the reach and redirected its arm after the visual cue appeared, resulting in longer RTs and non-predictive correct responses. Thus, movements that had previously been generated predictively from memory became more dependent on the visual instruction cue after anisomycin injection.

In this representative session, the effect of anisomycin was more pronounced for the short-term-trained sequence than for the more extensively trained sequences. The increase in error rate was significantly greater for the short-term-trained sequence than for the long-term- and extended-trained sequences (Fig. S1N; GLM, *P* < 0.001). Similarly, the decrease in predictive responses and the increase in RT were significantly greater for the short-term-trained sequence than for the extended-trained sequence (Fig. S1O, P; GLM for predictive responses and two-way ANOVA for RT, *P* < 0.001; see Materials and Methods).

In this representative session, performance of all three Repeating sequences deteriorated after injection (Fig. 3). The percentages of correct and predictive responses decreased significantly, whereas RT increased significantly (*χ² tests* and *t-tests*, *P* < 0.05). After anisomycin injection, we continued daily training and monitored task performance. We observed no overt behavioral abnormalities outside the task or other apparent side effects. With continued training, performance recovered toward pre-injection levels over the next 2–3 days (Fig. 3). We found no significant difference in recovery time among the three sequences. Because the anisomycin effect and recovery followed a stable time course (see also Fig. S13), we could confidently perform successive injections at appropriate time points. The 2–3-day recovery observed in this representative session is consistent with previous rodent studies reporting suppression of protein synthesis for approximately 48 hours after local anisomycin injection (*35*).

### Control experiments distinguished the effects of protein synthesis inhibition from nonspecific M1 dysfunction

We used three complementary approaches to determine whether the behavioral deficits after anisomycin reflected protein synthesis inhibition rather than nonspecific suppression or dysfunction of M1.

First, we compared the anisomycin phenotype with the effects of direct M1 inactivation. We injected muscimol, a GABA_A_ receptor agonist, into the shoulder representation of M1 (Fig. S5; *n=2* injection sessions in Monkey S) and tested performance of the Random and Repeating tasks (Fig. 4A–D; Fig. S6). Anisomycin injections largely spared Random task performance. In contrast, muscimol impaired both Random and Repeating task performance. In this session, the largest effects were observed for movements directed toward target 1, including movement ‘3-1’, and RT was prolonged in both tasks (Fig. 4A–D; Fig. S6; *χ2 test* for error rate*, t-tests* for RTs*, P* < 0.05). The increase in error rate varied across movements (Random: ‘3-1’, 44.6%; ‘5-3’, 1.8%; ‘1-5’, 17.6%; Repeating ‘3-1’, 19.2%; ‘5-3’, 4.3%; ‘1-5’, -2.8%). For the most strongly affected movement ‘3-1’, reaches frequently undershot the target after muscimol injection, producing prominent amplitude errors in both tasks (41% of ‘3-1’ trials in the Random task and 32% of ‘3-1’ trials in the Repeating task; Fig. 4A, B, Fig. S6). Thus, direct suppression of M1 activity produced a movement selective motor deficit affecting visually guided as well as memory-guided reaching, a pattern distinct from the anisomycin-induced impairment.

**Fig. 4.**
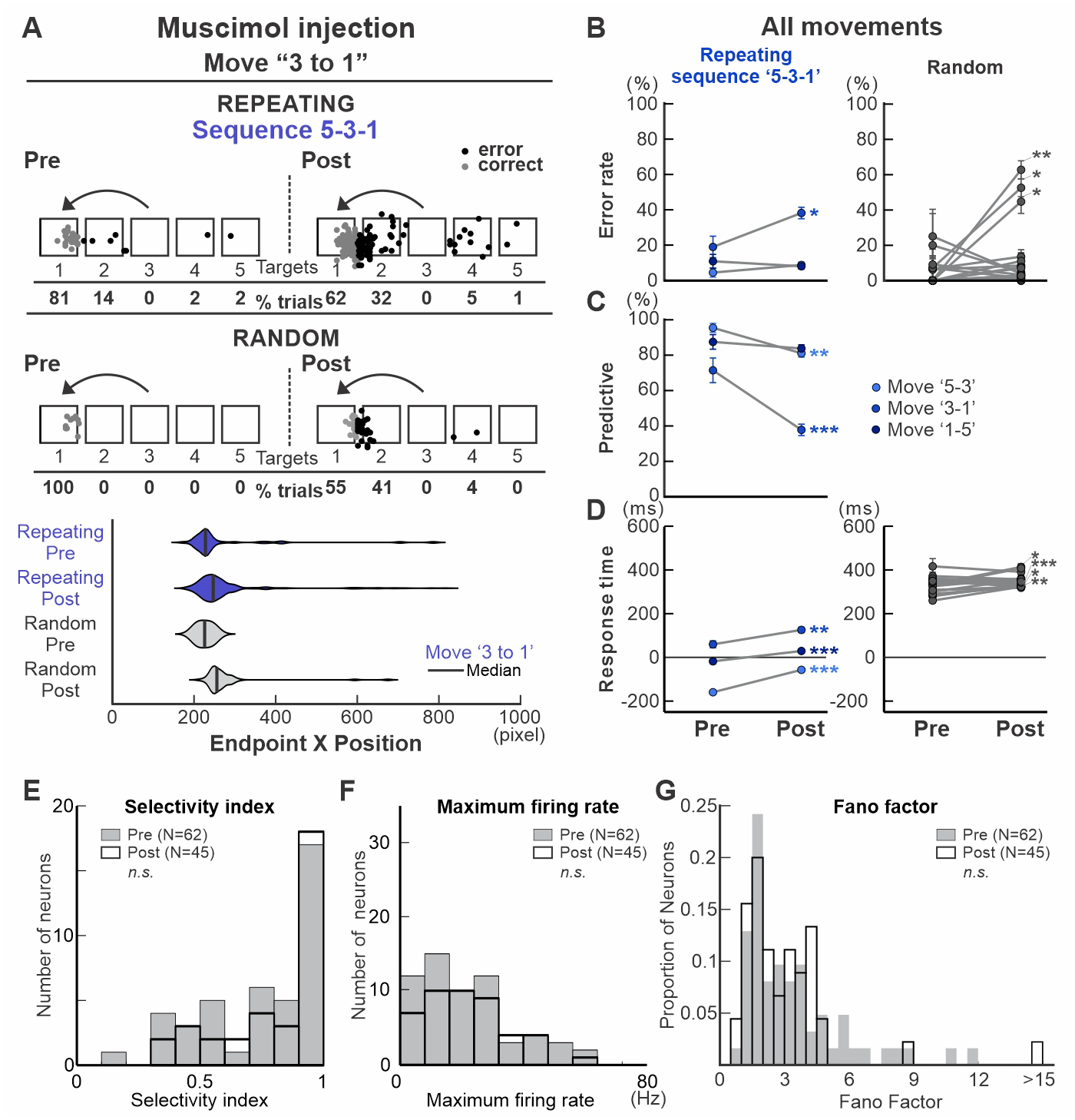
Control experiments examining movement execution and M1 neuronal activity. **(A–D)** Effects of muscimol injection into M1 on Repeating- and Random-task performance (Monkey S). **(A)** Reach endpoints for movement 3-1 of Repeating sequence 5-3-1 (top) and the corresponding movement in the Random task (middle), before and after injection. Gray and black dots indicate correct and error responses, respectively; percentages of responses to each target are shown below. Bottom, endpoint-position distributions for movement 3-1 in the Repeating and Random tasks before and after muscimol injection; vertical lines indicate medians. **(B–D)** Pre- and post-injection error rate (**B**), predictive-response rate (**C**), and response time (RT; **D**) for all component movements. Repeating-task movements are shown in blue and Random-task movements in gray. Pre–post differences were tested separately for each movement using Fisher’s exact test for error and predictive rates and Welch’s two-sample *t*-test for RT, with Holm correction across movements separately for each metric and task. Asterisks indicate Holm-adjusted significance (\**P* < 0.05, \*\**P* < 0.01, \*\*\**P* < 0.001). **(E–G)** M1 neural activity before and 24 h after anisomycin injection in Monkey R. **(E)** Distribution of movement-selectivity indices before (*n* = 62 neurons) and after (*n* = 45) injection; no significant difference was observed (two-sample *t*-test, *P* = 0.22). **(F)** Distribution of maximum firing rates, which also did not differ between pre-and post-injection recordings (two-sample *t*-test, *P* = 0.96). **(G)** Distribution of Fano factors before and after injection; spiking variability was unchanged (Wilcoxon rank-sum test, *P* = 0.3960).

Second, we tested whether anisomycin produced a loss of movement-related activity near the injection sites in M1. We recorded activity of neurons during the Random task before and after anisomycin injections, sampling neurons within 2 mm of the injection sites, inside the estimated spread of the drug. Representative neurons recorded before and after injection retained clear movement selectivity and comparable maximum firing rates during their preferred movements (Fig. S7). At the population level, we analyzed activity of 62 neurons recorded before injection and 45 neurons recorded after injection. We found no significant differences in movement selectivity, maximum firing rate, or trial-to-trial firing variability, quantified by the Fano factor between the pre- and post-injection populations (Fig. 4E-G; *P* > 0.05). Thus, anisomycin did not produce a detectable global suppression or loss of movement-related M1 activity at the time when behavioral impairments were present.

Third, we tested whether another type of protein synthesis inhibitor, cycloheximide, produced a behavioral effect similar to anisomycin. We injected cycloheximide into M1 and again observed impaired Repeating task performance, whereas Random task performance remained largely unchanged (Fig. S8). Cycloheximide injection increased error responses and reduced predictive responses in the Repeating task, although the magnitude of these effects varied across sequences and component movements. Because anisomycin and cycloheximide inhibit protein synthesis through distinct molecular mechanisms, their convergent behavioral effects support protein synthesis inhibition as the common mechanism underlying the disruption of learned sequence performance (*34*).

Together, these control experiments argue against a nonspecific loss of M1 function as an explanation for the anisomycin-induced behavioral deficit. Instead, they support the conclusion that the deficit of Repeating task performance following anisomycin injection reflects a specific consequence of induced translational blockade in M1 (*38, 44*), rather than nonspecific suppression of neural activity or motor function.

### Protein synthesis inhibition in M1 consistently impaired memory-guided sequence performance

To determine whether the behavioral effects observed in the representative experiment generalized across repeated injections, we analyzed all anisomycin-injection sessions and all available component movements in both monkeys (Monkey N, *n* = 5 sessions; Monkey R, *n* = 4 sessions; Fig. 5A, B; Figs. S9–S13). Injection sessions were separated by more than 2 weeks of training, and injections were placed at slightly different sites within the shoulder or elbow representations of M1 to minimize repeated tissue damage (see Materials and Methods).

**Fig. 5.**
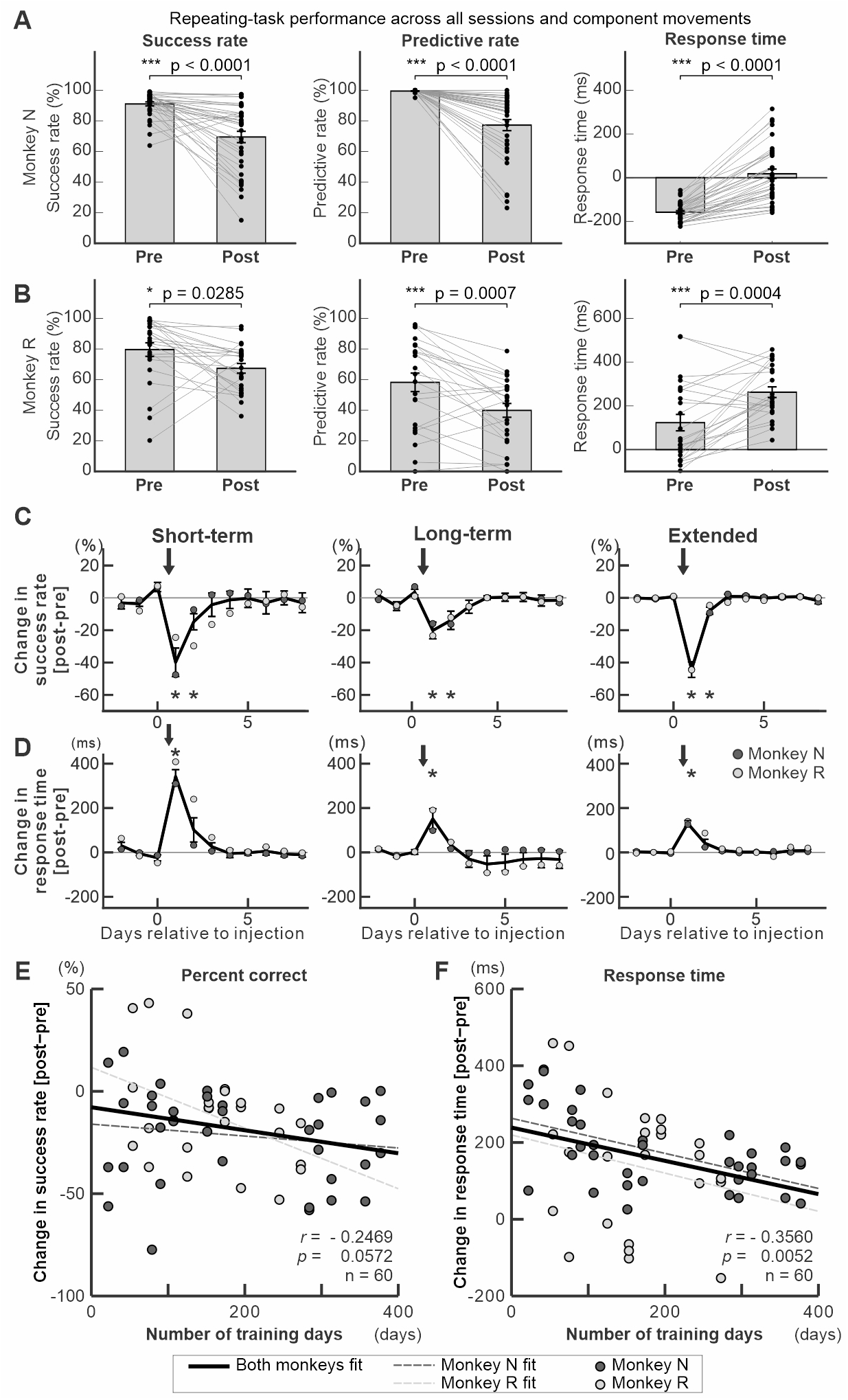
Population-level effects of anisomycin on Repeating-task performance and their relationship to training duration. **(A, B)** Pre- and post-injection performance across all anisomycin-injection sessions and all available component movements in Monkey N (**A**) and Monkey R (**B**). Left, success rate; middle, predictive-response rate; right, response time (RT). Each connected pair represents one component movement from one injection session. Bars show mean ± SE. Anisomycin significantly reduced success rate and predictive responses and increased RT in both monkeys. Exact *P* values are shown above each panel. **(C, D)** Recovery of anisomycin-induced changes in success rate (**C**) and RT (**D**) across short-term-, long-term-, and extended-training conditions. Values are expressed as post-minus-pre changes relative to the pre-injection day. Dark and light gray symbols denote Monkeys N and R, respectively; error bars indicate SEM. Arrows indicate anisomycin injection. Asterisks indicate significant differences from the last pre-injection value (*P* < 0.05). **(E, F)** Relationship between training duration and anisomycin effect size across all sessions and movements. Each symbol represents one movement from one Repeating sequence in one injection session. Black, dark-gray dashed, and light-gray dashed lines show ordinary least-squares fits for the combined dataset, Monkey N, and Monkey R, respectively. **(E)** Success-rate change showed a negative but nonsignificant relationship with training duration (mixed-effects slope = −0.0513 percentage points/day, *P* = 0.0829, 95% CI [−0.1095, 0.0069]). Pearson correlations were *r* = −0.2469 overall, −0.1573 in Monkey N, and −0.4221 in Monkey R. **(F)** RT change decreased significantly with training duration (mixed-effects slope = −0.4047 ms/day, *P* = 0.0054, 95% CI [−0.6850, −0.1244]). Pearson correlations were *r* = −0.3560 overall, −0.5720 in Monkey N, and −0.2192 in Monkey R. Mixed-effects models accounted for repeated observations within injection sessions.

Across the complete dataset, anisomycin significantly impaired Repeating-task performance in both monkeys. In Monkey N, anisomycin significantly reduced success rate and predictive responses and increased RT (all *P* < 0.0001; Fig. 5A). The same pattern was observed in Monkey R, with significant reductions in success rate (*P* = 0.0285) and predictive responses (*P* = 0.0007) and a significant increase in RT (*P* = 0.0004; Fig. 5B). Thus, the principal Repeating-task effects observed in the representative experiment replicated across injection sessions and animals, although the magnitude of the effect varied across component movements (Fig. S11). These movement-selective impairments also translated into deficits at the level of the complete three-movement sequence. Across sequence-level analyses, anisomycin reduced the probability of correctly completing the sequence and prolonged sequence completion time, with convergent evidence of impairment in both monkeys (Fig. S12).

In contrast, anisomycin did not significantly alter Random-task percent-correct performance or movement time (MT) in either monkey (Fig. S10, Monkey N: percent correct, *P* = 0.5702; MT, *P* = 0.2921; Monkey R: percent correct, *P* = 0.3208; MT, *P* = 0.3252). Random-task RT nevertheless increased significantly after anisomycin in both animals (Monkey N, *P* = 0.0002; Monkey R, *P* = 0.0192). Thus, visually guided reaching accuracy and movement execution remained largely preserved, despite some slowing of cue-guided response timing.

After each injection, performance recovered toward baseline over the following several days across sequences with different training histories (Fig. 5C, D; Fig. S13). This recovery was observed in both monkeys and across repeated injection sessions.

### The effect of protein synthesis inhibition in M1 decreased with extended practice

The representative experiment suggested that more recently trained sequences were more susceptible to anisomycin than extensively trained sequences (Fig. 3; Fig. S1N–P). We therefore asked whether this relationship generalized across the complete dataset. To avoid imposing arbitrary training-stage categories, we related the magnitude of the anisomycin effect for each sequence movement directly to the number of training days (Fig. 5E, F).

Because multiple movement observations were obtained within each injection session, we evaluated the relationship using linear mixed-effects models that accounted for repeated observations within sessions. The anisomycin-induced change in RT decreased significantly with training duration (slope = −0.4047 ms/day, *P* = 0.0054, 95% CI [−0.6850, −0.1244]; Fig. 5F). The reduction in success rate showed a similar negative relationship, but this effect did not reach statistical significance (slope = −0.0513 percentage points/day, *P* = 0.0829, 95% CI [−0.1095, 0.0069]; Fig. 5E). Descriptive Pearson correlations showed the same overall pattern for the combined dataset (RT: *r* = −0.3560, *P* = 0.0052; success rate: *r* = −0.2469, *P* = 0.0572). When examined separately for each monkey, RT showed a significant negative relationship in Monkey N (*r* = −0.5720, *P* = 0.00027) and the same negative directional trend in Monkey R (*r* = −0.2192, *P* = 0.3035). Success rate also showed negative correlations with training duration in both monkeys (Monkey N: *r* = −0.1573, *P* = 0.3596; Monkey R: *r* = −0.4221, *P* = 0.0399). Thus, the direction of the training-duration effect was consistent across monkeys for both RT and success rate, although statistical significance varied by metric and animal.

Together, these analyses show that the behavioral consequences of protein synthesis inhibition in M1 were greatest for more recently trained sequences and became smaller with extended practice, with the clearest effect observed for RT. Thus, the dependence of sequence performance on ongoing protein-synthesis in M1 declined as training progressed. This progressive reduction in anisomycin susceptibility supports the idea that memory-guided sequential movements undergo gradual stabilization in M1 over prolonged practice rather than becoming consolidated in a single discrete stage.

### Temporal evolution of RT impairment after anisomycin injection

To gain further insight into the processes underlying protein-synthesis-dependent consolidation of skilled sequential movements, we examined whether anisomycin-induced RT impairment changed over the course of individual sessions. We analyzed trial-by-trial RT changes before and after anisomycin injection (Figs. S14–S16). In the representative session shown in Fig. 3, RTs increased over the course of the post-injection session for several Repeating-task movements, whereas time or trial-dependent increases were weak or absent in the session before injection (Fig. S16A, B). Although the magnitude and direction of these within-session changes varied across individual movements and sessions, analysis of the complete nine-session dataset showed that within-session RT slopes were significantly more positive after anisomycin than before injection (trial-level linear mixed-effects model: β = 46.8 ms, 95% CI = 28.5–65.0 ms, *P* < 0.0001; complementary session-level paired analysis, *P* = 0.032; Fig. S16C). Thus, the RT impairment tended to develop further during post-injection task performance, although substantial heterogeneity was present across movements and sessions.

In two additional experiments, the monkeys were not trained for 3 or 5 days after anisomycin injection and were tested only after this delay. Under these conditions, we did not detect a significant impairment in either the Random or Repeating task when testing resumed (Figs. S15C, D and S16D; *P* > 0.05). Given the limited number of delayed-test experiments, these observations are preliminary but are consistent with the possibility that task engagement during the period of protein synthesis inhibition contributes to the subsequent behavioral effect.

## Discussion

In this study, we trained monkeys to learn sequential movements and interfered with information storage in the primary motor cortex (M1) at multiple time points during learning by injecting a protein synthesis inhibitor, anisomycin, into the arm area of M1. The anisomycin injection disrupted memory-guided sequential reaching at all stages of learning while largely preserving visually guided reaching accuracy and movement execution. Notably, the effect of anisomycin injection on sequence performance was more pronounced for the ‘Short-term’ trained sequence compared to the ‘Long-term trained’ and ‘Extended’ trained sequences. Further analysis demonstrated that the magnitude of the injection effect on the RTs of each movement was significantly correlated with the training duration of a sequence. These results suggest that the neural representation of ’Long-term’ and ‘Extended’ trained sequence is less susceptible to protein synthesis inhibition and thus less vulnerable to interference. The decreased susceptibility to protein synthesis inhibition suggests that the neural traces supporting sequential movements in M1 become gradually stabilized through repetitive practice.

Growing evidence demonstrates that M1’s activity changes with the learning of sequential movements both in the short term (within a training session to several months) (*1–7, 9–12*) and with years of training in humans and non-human primates (*7, 8, 13–29*). The changes in M1 after years of sequence training were reported in studies comparing musicians to non-musicians or comparing monkeys performing extensively trained sequential movements to those performing visually guided movements. However, it remains unclear from this work how learning-related plasticity occurs in M1, whether M1 is critically involved in sequence learning, and in which learning stage it may be. Changes of M1 activity after extensive training could be the outcome of plasticity during the early phase of the training. Our study using continuous long-term training and the staggered introduction of sequences covered learning stages within individual monkeys. Our results demonstrated that the protein synthesis inhibition disrupted the performance of memory-guided sequential reaching tasks at all stages of learning examined. This suggests that plasticity in M1 occurs at all stages of learning during repetitive practice over an extended period, continuously supporting learning of sequential movements.

The robust relationship between training duration and the RT effect suggests that extended practice progressively stabilizes processes in M1 that support efficient transitions between movements. The lack of a significant relationship between training duration and error-rate changes suggests that factors other than sequence age may also influence behavioral accuracy. One possibility is that other regions involved in sequence learning, such as premotor areas, compensate for M1 dysfunction differently across learning stages (*4, 45*). Additional sources of variability may include fluctuations in motivation, session length, fatigue, differences in movement distance or variation in the likelihood of direction errors between sequences, and the monkey’s ability to delay responses and wait for visual cues during the Repeating task. Together, these factors may contribute to the greater variability observed in error-rate measures.

The supplementary motor area (SMA) has long been considered responsible for the execution and maintenance of learned sequential movements (*4, 7, 46–50*). SMA neurons exhibit sequence specific activity (*49, 51*). Inactivation or lesion of the SMA disrupted the performance of memorized sequential movements, and inactivation or lesion of the pre-SMA disrupted the learning of a new sequence structure (*47, 49, 52–54*). In this view, M1 has been considered to relay the signal from the SMA to generate muscle commands for sequential movements. Additionally, we previously reversibly inactivated the dorsal premotor cortex (PMd) during performance of the same sequential movement tasks (*45*). PMd is densely interconnected with M1 and provides major cortical input to M1 (*55*). PMd inactivation selectively impaired memory-guided sequential movements while sparing visually guided reaching, suggesting that PMd contributes to the internal generation of sequential motor behavior. We cannot rule out the possibility that the SMA or PMd input to M1 (*55–57*) drives learning-related changes in M1. However, it is unlikely that performance disruption caused by anisomycin was the direct result of the drug’s spread to the adjacent PMd. Previous studies in monkeys have demonstrated that local cortical injections of pharmacological agents have a limited functional spread. For example, infusion of 3 μl of muscimol into monkey cortex affected neural activity within an area approximately 2–3 mm in diameter (*54*). Similarly, lidocaine injections required substantially larger volumes (i.e., 7 µl) to influence tissue located 2 mm away from the injection site (*58*). Because our injection sites were located at least 2 mm from the M1–PMd border and injection volumes were less than 3 μl each site, the effective spread was expected to remain largely confined within M1. Our results showed that protein synthesis inhibition disrupted the sequence performance at all stages of learning tested and suggest that aspects of learned sequences undergo protein-synthesis-dependent stabilization in M1 across the learning stages examined. One possibility is that protein-synthesis-dependent plasticity in M1 enhances synaptic efficacy and thereby contributes to increasingly fluent transitions between movements and the execution of individual movement components within a sequence. The movement-specificity of the effects in M1, while compatible with this area’s physiology, indicate that M1 encodes low-level features of movement sequences such as transitions or timing (*59*). Nevertheless, the results suggest that these hierarchically low representations are essential for skilled performance. Further studies are needed to explore these possibilities.

Studies in rodents using protein synthesis inhibitors suggest that memory may be dynamically modified or reconsolidated upon retrieval (*36, 37, 39, 60–63*). Specifically, upon memory retrieval, synapses within the memory network are first destabilized by protein degradation during reactivation, followed by stabilization to update or strengthen the memory through protein synthesis (*36, 37, 39, 62*). Supporting this view, the inhibition of protein degradation by *clasto*-lactacystin-*β*-lactone (*β*lac) has been shown to prevent anisomycin-induced memory impairment (*64*). Moreover, human studies using TMS and functional imaging suggest that memory reconsolidation may occur in humans as well (*65–67*). Several factors, including the age and strength of a memory (*60, 68, 69*), the novelty of the experience (*70, 71*), and duration of memory reactivation necessary for reconsolidation to occur (*68, 72*) are proposed as boundary conditions, which are considered to influence whether memory reconsolidation occurs.

Trial-by-trial analyses provided evidence that continued post-injection task performance enhanced the anisomycin-induced slowing of performance (Fig. S14, S16). In the representative experiment, RT increased progressively during the post-injection session. Similar patterns were observed in other sessions, although this effect was not consistent across the complete dataset. In two additional experiments in which the monkeys were not trained for 3 or 5 days after anisomycin injection, no significant impairment was detected when testing resumed (Fig. S15). Although these observations are limited, they are consistent with the possibility that task engagement during the period of protein synthesis inhibition contributes to the behavioral effect. This possibility is compatible with studies in rodents proposing that memory reactivation can trigger protein-synthesis-dependent restabilization or reconsolidation (*36, 37, 39, 60–63*).

Previous studies have shown that memory can be updated through reconsolidation when new information needs to be integrated (*73*), and that both the age and strength of a memory influence whether reconsolidation occurs or not (*68, 72*). Specifically, older and stronger memories (i.e., those reinforced through extended training) tend to be less sensitive to protein synthesis inhibition. In our study, susceptibility to anisomycin decreased with training duration, most clearly for RT (Fig. 5F). In our task, performance fluctuations due to movement variability may represent a form of “novelty” that serves as a boundary condition for reconsolidation. Our findings raise the possibility that repeated practice engages cycles of reactivation and protein-synthesis-dependent restabilization in M1 rather than producing plasticity solely through continuous incremental accumulation. This is consistent with the ‘lingering consolidation’ hypothesis, in which ‘the reactivation and reconsolidation cycle progressively stabilizes a memory’ (*37*). Furthermore, this would align with the gradual and continuous improvement of motor skill performance observed during extensive, repetitive practice. Recent studies in rodents have suggested that the motor cortex of rodents may not be essential for maintaining certain types of motor skills after “long-term” training (*74, 75*). In contrast, our results demonstrate that protein synthesis inhibition in M1 of monkeys significantly impaired sequence performance even after more than 300 days of training (Fig. 5). These differences may reflect several factors, including differences in whether the animals’ performance reached the boundary conditions of memory updating, the higher complexity of the sequential task used in our study, and species-specific anatomical differences. Further investigation will be needed to elucidate these possibilities.

Two-photon imaging studies offer insights into the possible mechanisms underlying memory-update in the motor cortex. In rodents, two photon imaging of the motor cortex during motor skill learning has shown the formation and growth of new dendritic spines and the elimination of old spines (*76, 77*). Additionally, two-photon imaging of cortical slices and electron microscopic studies revealed that protein synthesis is required for long-lasting synaptic plasticity (*38*) and spine-head enlargement and growth during learning (*78, 79*). The inhibition of protein synthesis resulted in a significant reduction in synapse number and synapse size in motor cortex of rodents in vivo (*34*). These observations suggest that, during motor skill training, unnecessary spines are eliminated through protein degradation, while the formation and growth of new spines are supported by protein synthesis which leads to enhanced neural efficacy. Thus, deficits in spine growth and formation may be associated with impairments in memory updates (i.e., consolidation), as observed in behavioral tests. Together with our findings, these studies provide a potential cellular framework in which repeated protein-synthesis-dependent synaptic modification contributes to the gradual stabilization of skilled motor performance (*36, 37, 39*).

Our findings suggest that M1 contributes to the gradual consolidation of skilled sequential movements across extended practice and that sequence performance becomes progressively less susceptible to protein synthesis inhibition as learning proceeds. This pattern is consistent with a lingering consolidation process in which memory stabilization gradually approaches an asymptote as a practiced motor memory becomes older and more stable. Nevertheless, our results indicate that protein-synthesis-dependent processes in M1 remain relevant even after more than 350 training sessions. These findings suggest that M1 makes a sustained contribution to the development and refinement of motor expertise, as part of a broader network through which repeated practice leads to increasingly skilled performance.

## Materials and Methods

### Behavioral task

We trained three monkeys (*Cebus apella*, male or female, weighing 1∼5 kg; > 2 years old) on the Random and Repeating tasks (Fig. 1) (*28, 29, 40, 45*). Each monkey was required to make sequential reaching movements to targets on a touch sensitive monitor with their right arms. In the Random task, the reaching movements were guided by visual targets displayed on a touch sensitive monitor in a pseudo-random order. Contact of the correct target triggered display of the next target after a 100 ms delay (Fig. 1B, D). After the monkey became proficient in performing the Random task (∼50 days of practice), we introduced the Repeating task. In the Repeating task, new targets were presented according to a repeating sequence of 3 elements (e.g., 5-3-1-5-3-1-…) 400 msec after contact of the preceding target (Fig. 1A, C, D). If the monkey made a correct response during the delay, the target is not shown, and the task increments to the next target in the sequence (Fig. 1A right panel, D middle panel). In both the Random and Repeating tasks, incorrect responses were followed immediately by illumination of the correct target and presentation of an error-feedback tone. The monkey was required to touch the correct target before the task could proceed. Thus, explicit feedback was available after every error, allowing the monkey to correct its response and resume the sequence from the appropriate position. With practice, a monkey memorized and performed the sequence of the Repeating task using predictive responses instead of visual cues to direct their movements in more than 80% of trials. The two tasks were performed continuously in alternating blocks of 200-500 trials (Fig. 1E). The monkey received a liquid reward after every 4-5 correct responses. After the monkey became proficient in the performance of the first sequence, additional sequences (e.g., 1-2-4-1-2-4…) were introduced (Fig. 1F, G). The new sequences were introduced in a staggered design with 50-200 days intervals. This training design enabled us to compare the effect of pharmacological injection between the ‘Long-term’ trained and ‘Short-term’ trained sequences in a single injection session. Each monkey learned two to three sequences (Monkey N: ‘5-3-1’, ‘1-2-4’, ‘2-3-4’; Monkey R: ‘1-2-4’, ‘2-3-4’; Monkey S: ‘5-3-1’). When the monkey frequently stopped working during a training session, the session was omitted and the behavioral data were excluded from further analysis because an insufficient number of trials and conditions were collected. Importantly, these exclusions were not related to poor learning or poor task performance. In most cases, the task was terminated during the Random-task block or occasionally at the beginning of the first Repeating-task block, typically after fewer than 100 trials and with a large number of no-response trials. Consequently, the monkeys performed little or no Repeating-task practice during these sessions, and there were insufficient data to evaluate sequential performance. Control of the behavioral task and collection of movement data including videos of task performance were performed by PC computers running TEMPO software (Reflective Computing, Olympia, WA) and Matlab. Touch screen signals, task related events and neural activity were collected on-line at 1 kHz.

### Behavioral estimates of learning phases

We used "Response Time" (RT) as the principal measure of sequence learning. We defined RT during the Random task as the time between the presentation of a new target and contact of that target. We defined RT during the Repeating task as the time between two targets touches minus the delay time, 400 msec (*28, 29, 40, 45*). We subtracted 400 ms to account for the delay in the cue presentation (Fig. 1D). This could result in a negative RT if the monkey moved quickly to the next target in the sequence before the presentation of a cue. RTs less than 150 ms were considered to be predictive. RTs less than 150 ms were chosen as a conservative cut-off for predictive responses as it is too fast for a simple reaction time to the visual cue. During the learning, an animal’s RT became shorter with practice (Fig. 1F, G). The data collected during the post-injection recovery period (1–3 days after injection) were not included in Fig. 1F and G. These figures were intended to illustrate the time course of learning under normal training conditions. Therefore, only data from training sessions without pharmacological injections were included. Inclusion of the post-injection data would have introduced transient injection-related effects and obscured the gradual learning-related changes in response time. A negative RT indicates that the animal was internally generating the response before the visual cue was presented. After about 50 days of training on the Repeating sequence, the monkeys made predictive responses (RT < 150 ms) without using visual cues in more than 80% of the trials (Fig. 1F, G).

#### Surgical Procedures

All experimental procedures were conducted according to NIH guidelines and were approved by the Institutional Animal Care and Use Committee (IACUC) of the University of Pittsburgh (protocol #23091930). We implanted a head restraint device, along with a MR compatible chamber for micro-injections and neural recording, on an animal’s skull using small screws and dental acrylic (*40, 45*). All surgical procedures were performed under general anesthesia using aseptic techniques. Anesthesia was induced with ketamine (10 mg/kg, IM) and maintained to surgical levels with 1-2.5% isoflurane. The animal received fluids throughout the surgical procedure (6-10 cc/hr, IV) and glycopyrolate (0.01 mg/kg, IM) to reduce secretions. Body temperature was maintained at 37-38 °C with a heating pad. We monitored body temperature, breathing, heart rate, end-tidal CO_2_, and O_2_ saturation. After the surgery, the animal received broad spectrum antibiotics (ceftriaxone, 75 mg/kg/day, IM, ∼5 days) and analgesics (buprenorphine, 0.01 mg/kg, IM, ∼3 days). The chamber placement over M1 in the left hemisphere was verified using structural MRI scans acquired before and after surgery (Fig. 2B). MR procedures for pretreatment, anesthesia, monitoring of vital signs and post-anesthesia monitoring were identical to surgical procedures, except that analgesic and antibiotic were not be given. After more than 2 weeks of recovery, we accustomed the animal to perform the tasks with its head restrained. When task performance returned to the pre-surgical level, we performed a craniotomy to expose the dura matter overlying M1 in the chamber using the same anesthetic techniques and safeguards used in the implant surgery. Animals were given dexamethasone (0.5 mg/kg divided into 3 doses, IM) on the day of the craniotomy surgery to prevent brain swelling.

#### Intracortical stimulation

We used intracortical microstimulation to identify the body part represented at each site in M1 and to physiologically define the border between M1 and the PMd (*40, 45, 55*). We used glass-coated micro-electrodes (0.6-2 MΩ at 1 kHz) to deliver intracortical microstimuli (*40, 45, 55*). A constant-current stimulator was used to deliver cathodal pulses (10–20 pulses, 0.2 ms duration, 333 Hz, 1-40 µA) at a depth of 1500 µm below the cortical surface. Stimulus intensity was measured with a current monitor (Ion Physics). The motor response evoked by stimulation was determined by visual observation and muscle palpation. The response threshold was defined as the lowest stimulus intensity necessary to evoke a response on more than 50% of the trials (*55*). We systematically mapped M1 with micro-electrode penetrations spaced ∼1.0 mm apart (except to avoid blood vessels) (Fig. 2).

#### Micro-injections

We made micro-injections of pharmacological agents (anisomycin, cycloheximide or muscimol) at selected sites in the forelimb area of M1 of a trained monkey (*40*). All injections were made into M1 of the left hemisphere, which was contralateral to the arm used to perform the task. We prepared solutions of anisomycin (100 µg/µl in ACSF, pH 7.2-7.4), cycloheximide (40 µg/µl in 20% ethanol/ACSF) and muscimol (5 mg/ml in saline) from commercially available powders (Sigma-Aldrich, MO). We injected the test solution at 1.5 mm below the cortical surface using a 30-gauge cannula connected to a 10 µl Hamilton syringe by applying slow pressure (e.g., over 10 minutes). Injection sites were placed more than 2 mm away from the border between M1 and PMd identified by microstimulation (Fig. 2). Previous studies using monkeys have shown that injection of 3 µl of muscimol into the cortex inhibited neural activity within a diameter of approximately 2-3 mm (*54*). Given that M1 in Cebus monkeys is more than 5 mm in width (Fig. 2), the effect of an injection of less than 3 µl of a pharmacological agent 2 mm away from the border was expected to remain confined within M1. The cannula was left in place for more than 5 min to allow diffusion of the solution and prevent its reflux. For each anisomycin injection session, we injected a total of 5 µl of anisomycin solution into M1 of two monkeys. The solution was divided into two aliquots (2.6 µl and 2.4 µl) and injected at sites within M1 where microstimulation elicited elbow or shoulder movements. The injection volume was divided to minimize potential tissue damage and to ensure broader coverage of the cortical regions representing the elbow and shoulder. Task performance was assessed 20–24 hours after the injection, within the drug’s active window (48-72 hours), and daily thereafter. For control experiments, task performance was first assessed outside of the drug’s active window, 3-5 days after the injection. Measurement of performance on days prior to an injection and/or on trial blocks preceding an injection was used as the baseline for comparison with post-injection performance. We repeated the anisomycin injections with behavioral testing five times in monkey N and four times in monkey R to test its effect on learning. To verify that the effect on behavior is due to the common effect of the agents (i.e., protein synthesis inhibition), we injected 3 µl of cycloheximide (40 µg/µl) at a site within M1 where microstimulation elicited elbow or shoulder movements. Cycloheximide blocks protein synthesis by a different mechanism than anisomycin. The effect of cycloheximide injection on tasks was tested 20-24 hours after the injection. Measurement of performance on a day prior to an injection and/or on trial blocks preceding an injection was used as the baseline for comparison with post-injection performance. Muscimol, a GABA_A_ agonist, (1-3 µl) was injected at a site within M1 of the monkey S where microstimulation elicited elbow or shoulder movements immediately after the collection of the baseline behavioral data and the effect of injection on task performance was tested 20 min after the injection to allow diffusion of the chemical in the brain tissue (*n*=2). After each injection session, the task performance was monitored every day until it recovered to the baseline to track the emergence of injection effects and to assess recovery from the drug’s effects. Injections were separated with more than 7 days and 5 training sessions. We examined whether the injection of the pharmacological agent impaired performance of the Random and Repeating tasks. During a post-injection test session, blocks of Random and Repeating trials alternated at frequent intervals to sample the animal’s performance evenly as the effect of the test substance emerged and intensified.

#### Analysis of performance

For every trial of each task session, we recorded various task parameters and measures of performance. The effect of an injection was assessed by examining the following: percentage of correct responses, types of incorrect responses, percentage of predictive responses (defined as a correct response with RT < 150 msec), Response Time (RT) and Movement Time (MT) (*40, 45*). The errors recorded were: no hit, wrong target hit, background hit, or a corrective response (a correct response that immediately follows an error). A trial was defined as the interval beginning when the monkey contacted a target and ending when the monkey contacted another target, or when the maximum response time (800 ms) elapsed. Therefore, each movement between two targets was considered one trial. No responses triggered the start of another trial and thus also incremented the trial count. MT is defined as the interval between the release of contact from one target to touch of the next target. We defined RT during the Random task as the time between the presentation of a new target and contact of that target. We defined RT during the Repeating task as the time between targets touches minus the delay time, 400 msec (see the section of Behavioral estimates of learning phases above) (*40, 45*). RT less than 150 ms was chosen as a cut-off for predictive responses as it is too fast for a simple reaction time to the visual cue. A wrong target hit during the Repeating task was categorized as two types: *amplitude errors* and *direction errors*. An *amplitude error* is defined as a reach performed in the correct direction, but to an endpoint outside of the correct target. This type of error suggests a deficit in motor production. A *direction error* was defined as a reach performed in the direction opposite to the correct target. This type of error suggests a deficit in selecting the movement component in the sequence. Direction errors could only occur for movements starting from targets on the center of the touch monitor (target 2, 3, 4), since movements in the opposite direction from targets at the edge of the touch monitor would fall outside the touch monitor. For statistical analysis, both types of errors were included in the calculation of performance error. To further characterize the deficits, we reclassified error responses according to their relationship to the learned sequence. Errors were categorized as Out-of-Sequence or Off-Phase. Out-of-Sequence errors occurred when the monkey reached to a target that was not part of the current sequence. Off-Phase errors occurred when the monkey reached to a target that belonged to the sequence but was selected at the incorrect phase of the sequence. Although both error types reflect deficits in target selection, they represent distinct error patterns. An increase in RT and decrease in the number of predictive responses suggest an increase in the time for movement selection. Corrective responses were removed from analysis because the target is predictable after an error. We also monitored movement kinematics during the task using high speed video recording (100 Hz, Basler Inc., PA) in the frontal plane. For statistical analysis, we used *χ^2^* tests with Holm–Bonferroni’s correction to examine the significance of changes in success rate and predictive responses in a session. We used *t-tests* with Holm–Bonferroni’s correction to examine MT and RT changes. For the summary plots of the representative sessions (Figs. S1, S6, S8-S9), trials from all three movements of each Repeating sequence, or from all movements of the Random task, were pooled to obtain pre- and post-injection estimates. Predictive response rate was calculated as the number of correct responses with RT < 150 ms divided by the total number of trials. Pre–post differences were tested using trial-level models that included injection condition and movement identity as predictors: binomial GLMs were used for error and predictive response rates, and linear models were used for RT and MT. Error bars indicate 95% Wilson confidence intervals for proportions and 95% confidence intervals for RT and MT means. To test the reproducibility of the task performance results, we repeated the injections five times in monkey N and four times in monkey R. In monkey R, we performed three additional injections to examine the effect of anisomycin injection on neural activity (see the Electrophysiology section of the Methods). To compare the strength of the injection effect between the sequences in the different learning phases, we fitted a generalized linear model (GLM), using either performance or predictive responses as the variable and the learning phase as a predictor for analysis shown in Fig. S1N-P. A binomial distribution with a logit link function was applied. We assessed the relationship between training duration and post-minus-pre changes in success rate and RT using *Pearson’s correlation coefficients*, including all component movements from all Repeating sequences and injection sessions. To account for repeated movement observations within injection sessions, we also fitted linear mixed-effects models with training duration as a fixed effect and injection session as a random effect; monkey identity was included as a fixed effect in the combined analysis. As a sensitivity analysis, we performed a session-cluster bootstrap by resampling entire injection sessions with replacement (10,000 iterations) and calculated 95% confidence intervals for the regression slope. For the within-session RT analysis (Fig. S16A–C), only correct first-attempt responses were included. The first 20 and last 20 responses for each movement were compared using two-sided Welch’s *t* tests, with Holm correction across the three movements within each sequence (Fig. S16A, B). RTs were also ordered chronologically within each movement and normalized from 0 to 1. In the representative session, within-session RT slopes were estimated separately for each movement by linear regression, and PRE–POST slope differences were tested by the response-order × condition interaction, with Holm correction within each sequence. Population-level differences across all nine anisomycin-injection sessions were tested using a linear mixed-effects model with response order, condition, their interaction, sequence, and monkey as fixed effects, random intercepts and response-order slopes for session, and a random intercept for session-by-movement. MATLAB (MathWorks) was used for statistical analyses.

#### Electrophysiological recording and analysis of neural activity

To evaluate the effect of pharmacological agents on neural activity, we recorded extracellular activity of single neurons in M1 using glass-coated Elgiloy electrodes (0.6-1.5 MΩ). We recorded neural activity before and after the injection of anisomycin while the monkey S performed the tasks. As shown in Fig. 3 and 5, an anisomycin injection disrupted performance of the Repeating task on up to 80% of trials. Therefore, to evaluate the effect of injection on neural activity, we limited analysis on the data collected during the Random task. Neural data after the anisomycin injection were obtained during four injection sessions. Of these, performance data of three injection sessions were excluded from behavioral analysis because of insufficient number of trials in the Repeating task. The sample of neurons was taken 20-24 hours after the injection within 1-2 mm of the injection site from regions of M1 where intracortical stimulation evoked shoulder, elbow or wrist movements at thresholds lower than 25 µA. For each neuron, we used maximum firing rate and a selectivity index (SI) to assess the effect of injections. For each Random move, we measured the mean firing rate of a neuron in a 200 msec interval centered on target contact or release. We used this value to calculate a SI for each neuron: SI = [Max -Min]/ [Max + Min]. For statistical analysis, we used *t-test* to examine changes in SI and max firing rate between the group of M1 neurons recorded before and after the injections. To quantify trial-by-trial spiking variability, the Fano factor (FF) was calculated for each isolated single neuron during a 200 msec window centered on target contact (*80*). For each neuron, the total spike count within the 200-ms epoch was computed for each individual trial. The Fano factor was defined as the ratio of the variance to the mean of these trial-by-trial spike counts (*FF* = σ^2^/µ). Neurons with a mean spike count of zero were excluded from further analysis to prevent division by zero. Changes in spiking variability between the pre-and post-injection populations were statistically evaluated using a two-tailed Wilcoxon rank-sum test (*p* = 0.05).

## Acknowledgments

We thank Dr. Peter L. Strick for support, discussions and suggestions; Dr. Floh Thiels for discussions; Moya Carrier for animal training and assistance; Mike Page for task programming.

## Author contributions

Writing (original draft, review & editing), Conceptualization: MO, NP Methodology, Investigation, Resources, Funding acquisition, Dara curation, Validation, Supervision, Dormal analysis, Software, Project administration, Visualization: MO

## Data, code, and materials availability statement

All data needed to evaluate the conclusions in the paper are present in the paper and the Supplementary Materials. The raw data and custom Matlab code used for data analysis and for evaluating and reproducing the study are publicly available in Zenodo (https://doi.org/10.5281/zenodo.23348172). The detailed task procedures and the detailed step-by-step surgical and chamber maintenance protocols are available at Bio-protocol (https://doi.org/10.21769/BioProtoc.3719) and protocols.io (https://dx.doi.org/10.17504/protocols.io.dm6gp4ro1gzp/v1), respectively. This study did not generate new materials.

