## Supplementary Materials for "Gradual consolidation of skilled sequential movements in primary motor cortex of non-human primates"

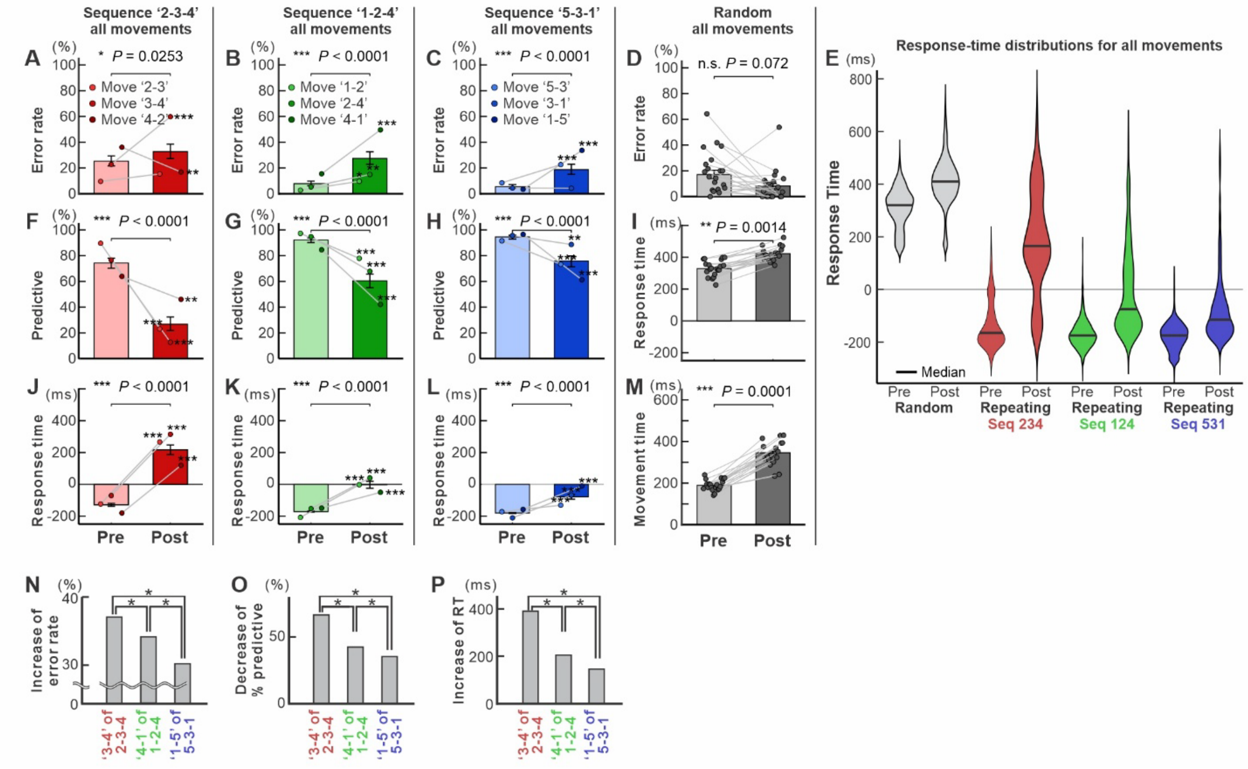


**Fig. S1. All-movement analysis and training-stage dependence of anisomycin effects in the representative session shown in Fig. 3.**

**(A–C, F-H, J-L)** Pre- and post-injection performance for all component movements of the three Repeating sequences: sequence '2-3-4' (**A, F, J**), sequence '1-2-4' (**B, G, K**), and sequence '5-3-1' (**C, H, L**). Error rate (**A-C**), predictive-response rate (**F-H**), and response time (RT; **J-L**) are shown. Bars represent trial-weighted pooled estimates across the three movements, and connected points show movement-specific estimates. Pre–post differences for individual movements were tested using Fisher's exact test for error and predictive rates and Welch's two-sample *t*-test for RT; *P* values were Holm-adjusted across the three movements separately for each metric. Asterisks adjacent to individual movements indicate Holm-adjusted significance (*$P < 0.05$, **$P < 0.01$, ***$P < 0.001$).

**(D, I, M)** Error rate (**D**), RT (**I**), and movement time (**M**) for all movements performed in the Random task. Connected points indicate the same movements before and after injection; bars show pooled estimates.

**(E)** RT distributions before and after injection for all movements in the Random task and each Repeating sequence. Horizontal lines indicate medians.

**(N–P)** Magnitude of the anisomycin effect for the representative movements shown in Fig. 3 across the short-term-, long-term-, and extended-trained sequences ('2-3-4', '1-2-4', '5-3-1', respectively): increase in error rate (**N**), decrease in predictive-response rate (**O**), and increase in RT (**P**). Asterisks indicate significant pairwise differences among training stages (*$P < 0.05$).

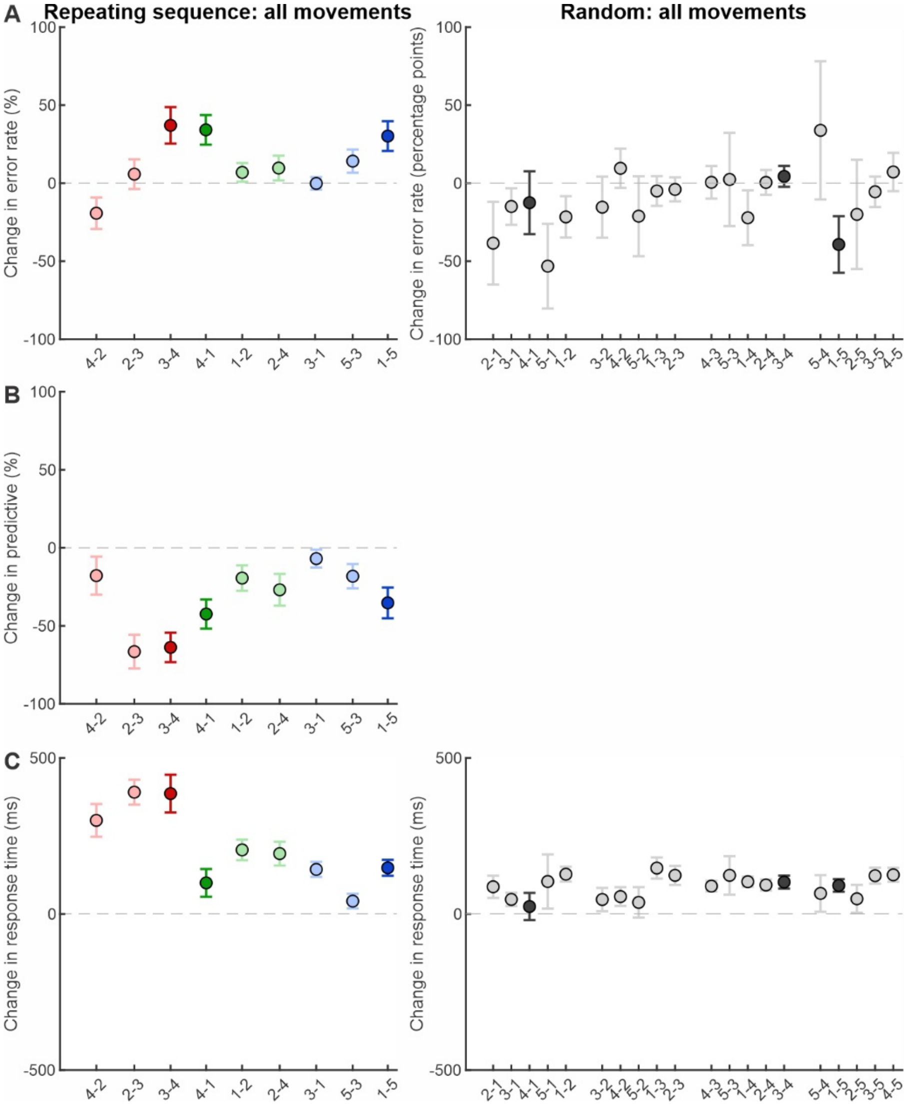


**Fig. S2. Movement-specific effects of anisomycin in the representative injection session shown in Fig. 3.**

Changes in performance from pre- to post-injection are shown separately for individual movements in the Repeating task (left) and Random task (right) of Monkey N, session A5. Values were calculated as post-injection minus pre-injection performance.

**(A)** Change in error rate. **(B)** Change in predictive-response rate. **(C)** Change in response time (RT). For the Repeating task, movements from sequences '2-3-4', '1-2-4', and '5-3-1' are shown in red, green, and blue, respectively. Darker symbols indicate the representative movements shown in Fig. 3 ('3-4', '4-1', and '1-5'), whereas lighter symbols indicate the other component movements within each sequence. For the Random task, all available movements are shown in gray; darker symbols indicate movements corresponding to the representative Repeating-task movements. Error bars indicate 95% confidence intervals. These plots illustrate the heterogeneity of the anisomycin effect across individual movements.

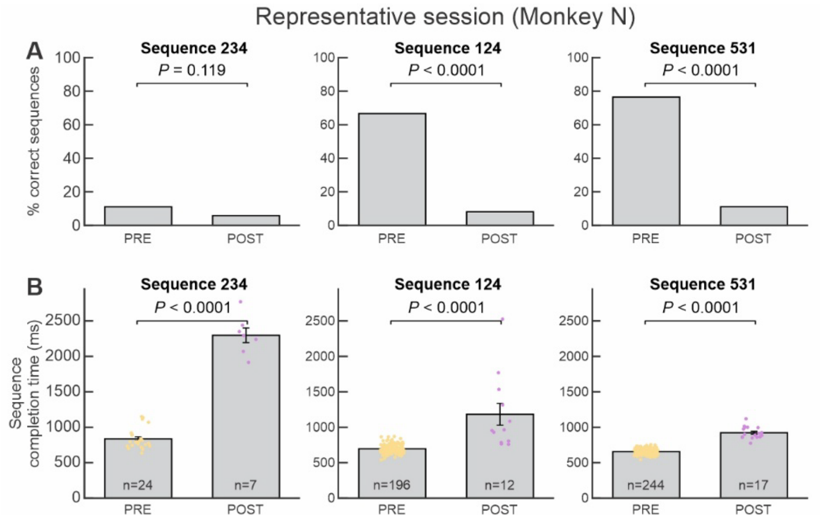


**Fig. S3. Effects of anisomycin on sequence-level performance.**

**(A, B)** Sequence performance in the representative Monkey N injection session A5 shown in Fig. 3, reanalyzed using each complete three-movement sequence as the unit of analysis. A sequence was scored as correct only when all three component movements were completed correctly without an intervening error; corrective responses following an error were not counted as successful completion of that sequence. Anisomycin markedly reduced complete-sequence accuracy for sequences '1-2-4' and '5-3-1' and significantly prolonged sequence completion time for all three sequences.

(**A)** Percentage of correctly completed sequences before (PRE) and after (POST) anisomycin injection for sequences '2-3-4', '1-2-4', and '5-3-1'. Statistical comparisons were made using Fisher's exact tests. The short-trained sequence '2-3-4', still at an early stage of learning, had relatively low pre-injection sequence accuracy, limiting the extent of a further decline in this measure.

(**B)** Sequence completion time for correctly completed sequences, defined as the elapsed time required to complete one full three-movement sequence cycle. Dots indicate individual correctly completed sequences; bars and error bars indicate mean ± SE. Sample sizes indicate the number of correctly completed sequences contributing to the completion-time analysis. Statistical comparisons were made using Wilcoxon rank-sum tests.

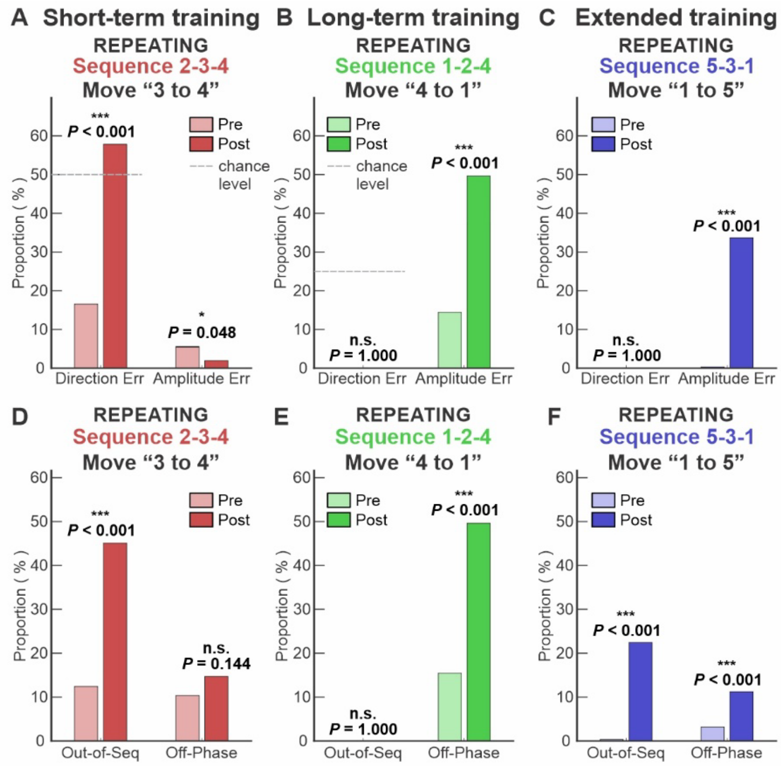


**Fig. S4. Error analysis for the representative anisomycin-injection session shown in Fig. 3.** **(A-C)** Errors were classified as Direction Errors or Amplitude Errors. Direction errors occurred when the monkey selected an incorrect movement direction (e.g., a movement in the direction opposite to the correct target). Amplitude errors occurred when the movement undershot or overshot the intended target. Therefore, direction errors occur for movements starting from the targets in the center of the screen (e.g., movements starting from targets 2, 3, and 4). For the '1-5' movement in sequence '5-3-1', direction errors could not occur because the starting position (target 1) was located at the edge of the monitor. Direction error increased significantly in sequence '2-3-4' (**A**), whereas amplitude error increased significantly in sequence '1-2-4' (**B**) and '5-3-1' (**C**) ($P < 0.001$). The chance level for Direction Errors is the proportion of targets located in the direction opposite to the correct target among the four possible target locations. For example, for the '3-4' movement in sequence '2-3-4', two of four possible targets were located in the direction opposite to the correct movement from target 3 to target 4 (i.e., left), resulting in a chance-level Direction error rate of 50%.

(**D-F**). In this analysis, we reclassified the error data shown in **A-C** based on target-selection errors. Errors were classified as Out-of-Sequence or Off-Phase. Out-of-Sequence errors occurred when the monkey reached to a target that was not part of the current sequence. Off-Phase errors occurred when the monkey reached a target that belonged to the sequence but was selected at the incorrect phase of the sequence. Both error types reflect deficits in target selection but represent distinct error patterns. Both Out-of-Sequences errors increased significantly in sequences '2-3-4' (**D**) and '5-3-1'(**F**). Off-Phase errors increased significantly in sequence '1-2-4' (**E**) and '5-3-1' (**F**) ($P < 0.001$). This session corresponds to injection A5 in Monkey N.

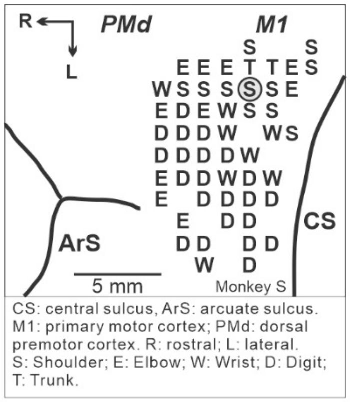


**Fig. S5. Intracortical microstimulation map and a muscimol injection site in M1 of Monkey S.**

Letters indicate the movements evoked by intracortical microstimulation at each site in M1. S: Shoulder; E: Elbow; W: Wrist; D: Digit; T: Trunk. Gray circle indicates a muscimol injection site. Monkey S was used exclusively for muscimol injections into M1. ArS: arcuate sulcus; CS: central sulcus; PMd: dorsal premotor cortex; M1: primary motor cortex. R: rostral; L: lateral.

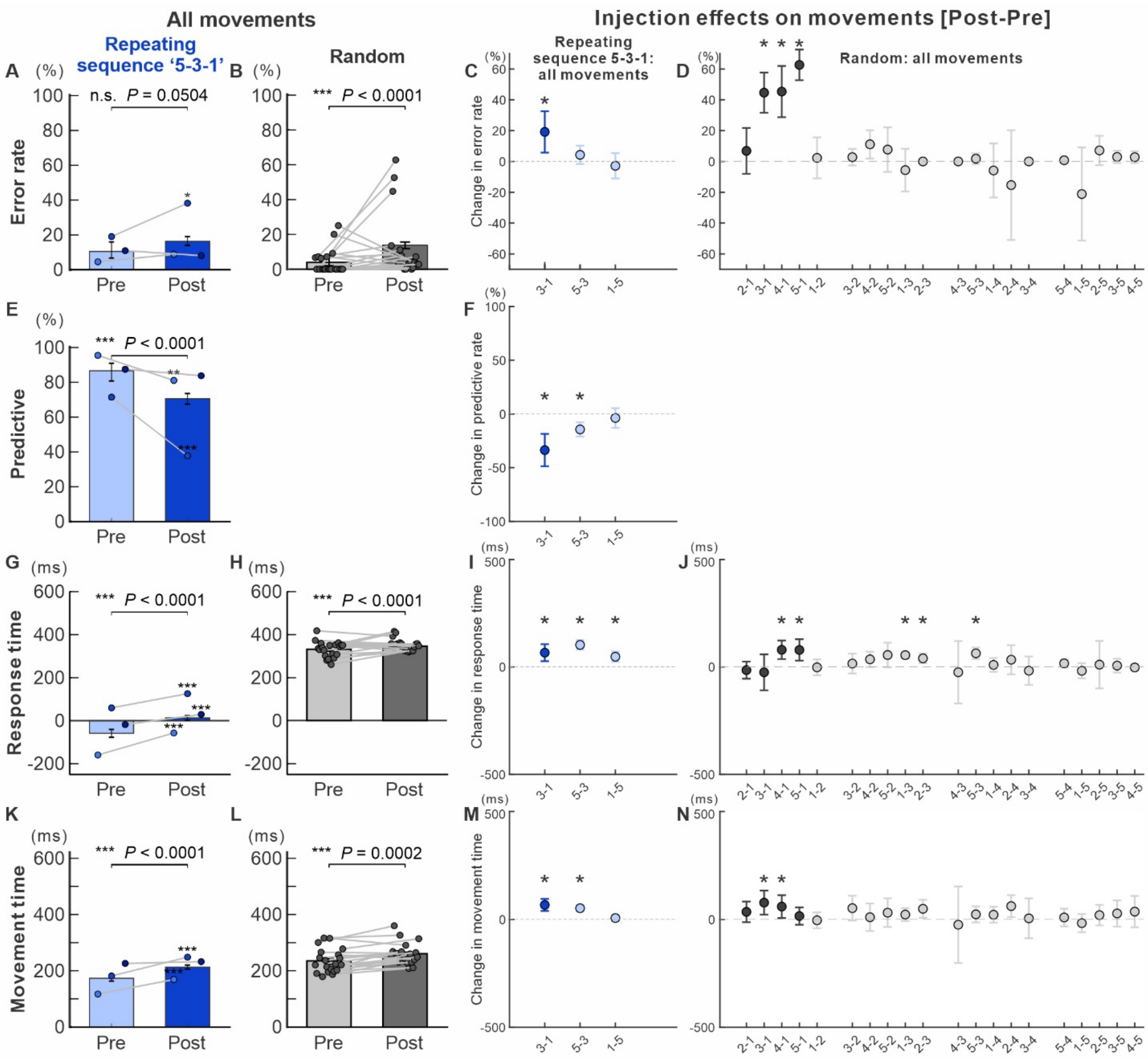


**Fig. S6. Movement-specific effects of muscimol on Repeating- and Random-task performance in Monkey S.**

**(A, E, G, K)** Repeating sequence '5-3-1'. Pre- and post-injection performance across all movements.

**(B, H, L)** Random task. Pre- and post-injection performance across all movements.

Error rate (**A, B**), predictive-response rate (**E**), response time (RT; **G, H**), and movement time (MT; **K, L**) are shown. Bars represent pooled estimates; error bars indicate 95% confidence intervals. Pre–post differences in the all-movement analyses were evaluated using trial-level models including injection condition and movement identity as predictors; exact *P* values are shown above the panels. Connected points show the corresponding movement-specific values. Asterisks indicate movement-specific significance (**P* < 0.05, ***P* < 0.01, ****P* < 0.001).

(**C, D, F, I, J, M, N)** Movement-specific injection effects, expressed as post-injection minus pre-injection values. (**C, F, I, M)** Component movements of Repeating sequence '5-3-1'; (**D, J, N)** all displayed movements in the Random task. Changes are shown for error rate (**C, D**), predictive-response rate (**F**), RT (**I, J**), and MT (**M, N**). For each movement, pre–post differences were tested using Fisher's exact test for error and predictive rates and Welch's two-sample *t*-test for RT and MT. *P* values were Holm-adjusted across movements separately for each metric and task. Asterisks indicate Holm-adjusted significance (*$P < 0.05$,). Error bars indicate 95% confidence intervals. These analyses illustrate the heterogeneity of the muscimol effect across individual movements.

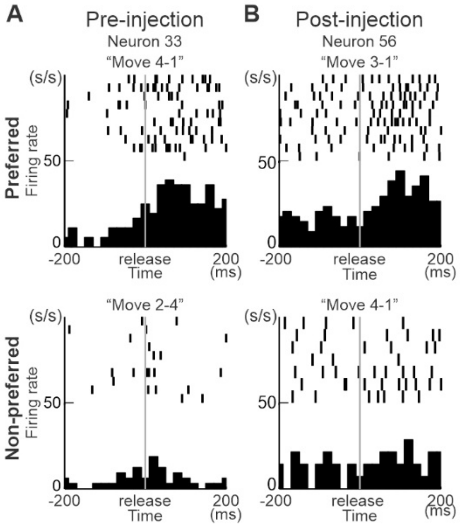


**Fig. S7. Representative M1 neural activity before and after anisomycin injection.**
**(A)** Activity of a representative movement-selective M1 neuron recorded before anisomycin injection.
**(B)** Activity of a representative movement-selective M1 neuron recorded after anisomycin injection. Raster plots and histograms are aligned to screen release (time 0). For each neuron, the upper panels show activity during the preferred movement, defined as the movement associated with the highest firing rate, and the lower panels show activity during a non-preferred movement associated with low firing. The preferred and non-preferred movements were '4-1' and '2-4', respectively, for the pre-injection neuron (Neuron 33), and '3-1' and '4-1' for the post-injection neuron (Neuron 56). Firing-rate histograms are shown in spikes/s. These examples illustrate movement-related activity patterns observed before and after anisomycin injection.

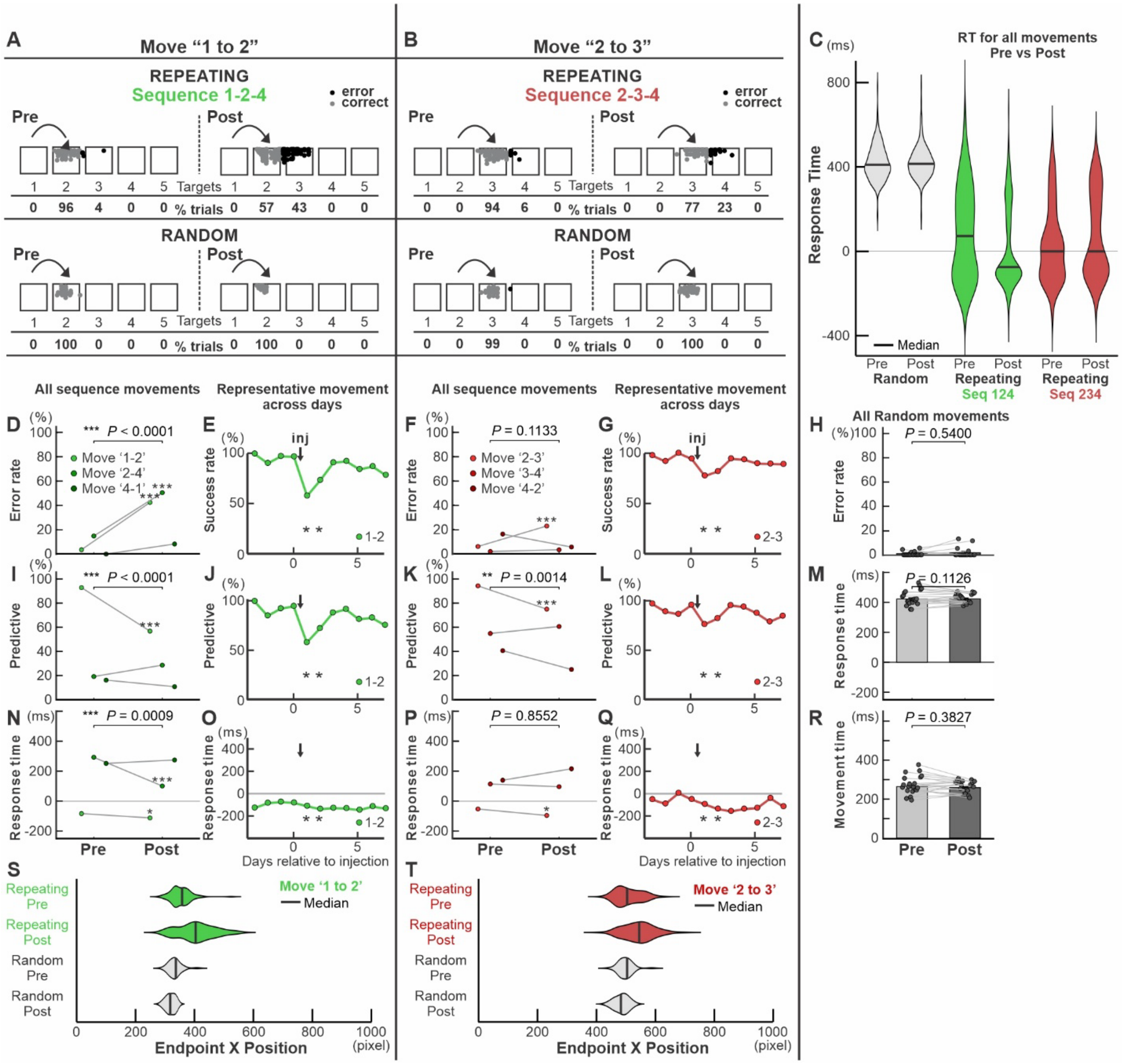


**Fig. S8. Effects of cycloheximide injection on Repeating- and Random-task performance in Monkey R.**

Cycloheximide was injected into M1 after the monkey had practiced sequence '1-2-4' for 340 days and sequence '2-3-4' for 460 days.

**(A, D, E, I, J, N, O, S)** Effects on sequence '1-2-4'.

**(A)** Reach endpoints for the representative movement '1-2' before and after cycloheximide injection during the Repeating task (top) and the corresponding movement in the Random task (bottom). Gray and black dots indicate correct and error responses, respectively; percentages of responses to each target are shown below.

**(D, I, N)** Pre- and post-injection error rate **(D)**, predictive-response rate **(I)**, and response time (RT) **(N)** for all three component movements of sequence '1-2-4'. Connected points show movement-specific values. Exact *P* values for the all-movement comparisons are shown above the panels. Asterisks adjacent to individual movements indicate significant pre–post changes after multiple-comparison correction (**P* < 0.05, ***P* < 0.01, ****P* < 0.001).

**(E, J, O)** Daily time course of success rate, predictive-response rate, and RT, respectively, for the representative movement '1-2'. Arrows indicate cycloheximide injection; asterisks indicate significant differences from the pre-injection baseline.
**(S)** Endpoint-position distributions for the representative movement '1-2' in the Repeating and Random tasks before and after injection; horizontal bars indicate medians.
**(B, F, G, K, L, P, Q, T)** Corresponding analyses for sequence '2-3-4', with movement '2-3' used as the representative movement.
**(C)** RT distributions before and after injection for all movements in the Random task and each Repeating sequence. Horizontal bars indicate medians.
**(H, M, R)** Random-task performance across all movements: error rate **(H)**, RT **(M)**, and movement time (MT) **(R)**. Connected points indicate corresponding movements before and after injection; bars show pooled estimates. *P* values are shown above the panels.
Cycloheximide impaired Repeating-task performance, although the magnitude and behavioral changes varied across component movements and sequences. In contrast, no significant changes were detected in Random-task error rate, RT, or MT.

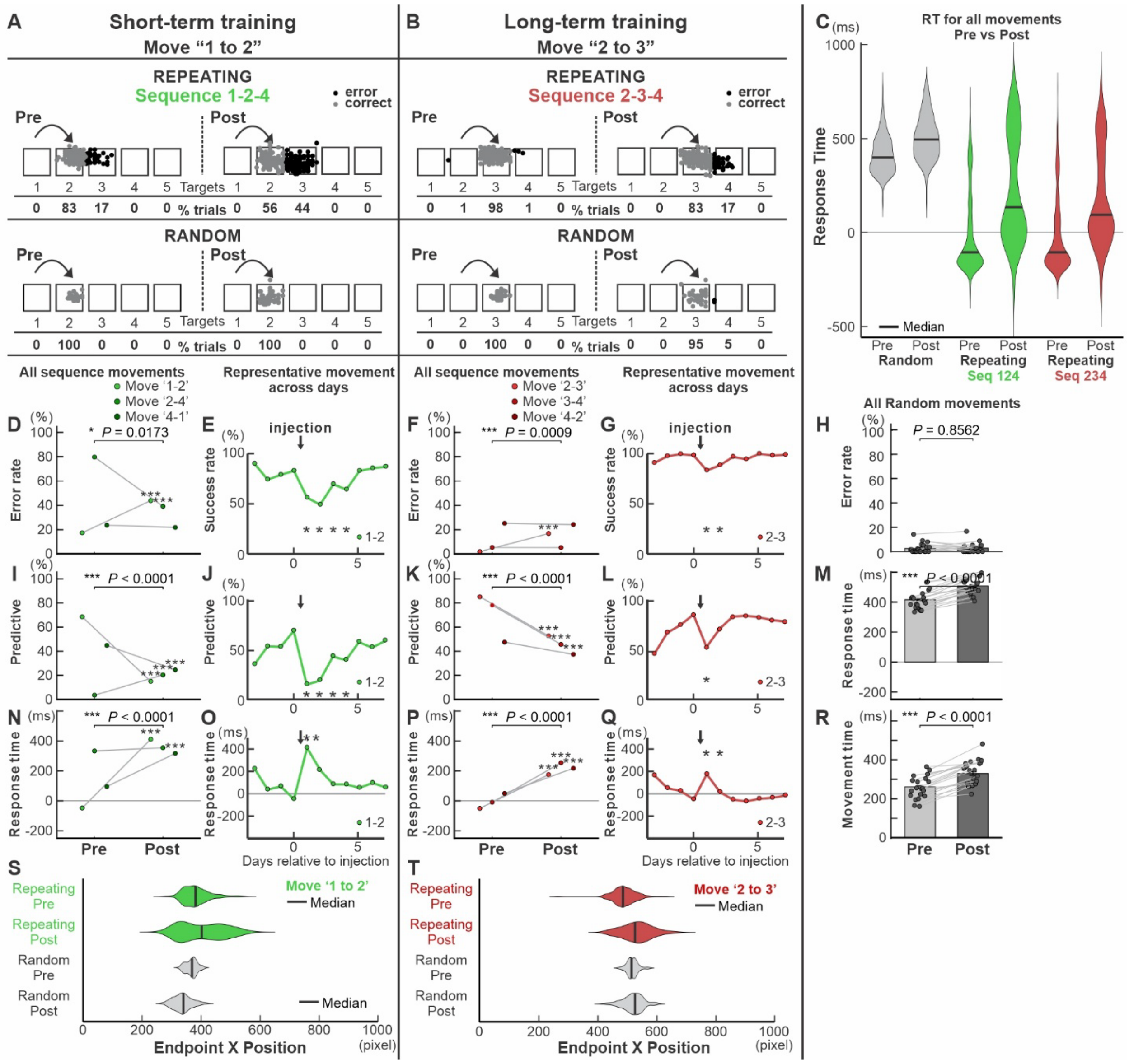


**Fig. S9. Representative effects of anisomycin on Repeating- and Random-task performance in Monkey R.**

Data are from anisomycin-injection session A1 of Monkey R. At the time of injection, sequence '1-2-4' had been trained for 54 days (short-term training) and sequence '2-3-4' for 174 days (long-term training).

**(A, D, E, I, J, N, O, S)** Short-term-trained sequence '1-2-4'.

(**A)** Reach endpoints for the representative movement '1-2' before and after anisomycin injection in the Repeating task (top) and the corresponding movement in the Random task (bottom). Gray and black dots indicate correct and error responses, respectively; percentages of responses to each target are shown below.

**(D, I, N)** Pre- and post-injection error rate **(D)**, predictive-response rate **(I)**, and response time (RT; **N)** for all three component movements of sequence '1-2-4'. Connected points show movement-specific values. Exact *P* values for the all-movement comparisons are shown above the panels. Pre–post differences for individual movements were tested using Fisher's exact test

for error and predictive rates and Welch's two-sample *t*-test for RT, with Holm correction across the three component movements separately for each metric. Asterisks adjacent to individual movements indicate Holm-adjusted significance (*$P < 0.05$, **$P < 0.01$, ***$P < 0.001$).
**(E, J, O)** Daily time course of success rate, predictive-response rate, and RT, respectively, for movement '1-2'; arrows indicate the injection day, and asterisks indicate significant differences from the pre-injection baseline.
**(S)** Endpoint-position distributions for movement '1-2' in the Repeating and Random tasks before and after injection; horizontal bars indicate medians.
**(B, F, G, K, L, P, Q, T)** Corresponding analyses for the long-term-trained sequence '2-3-4', with movement '2-3' used as the representative movement.
**(C)** RT distributions before and after injection for all movements in the Random task and in sequences 1-2-4 and 2-3-4; horizontal bars indicate medians.
**(H, M, R)** Random-task performance across all displayed movements: error rate (**H**), RT (**M**), and movement time (**R**). Connected points indicate corresponding movements before and after injection; bars show pooled estimates. Statistical comparisons were performed using the all-movement analyses described in Methods; exact *P* values are shown above the panels.
These data illustrate the overall impairment of Repeating-task performance and its heterogeneity across individual component movements in Monkey R.

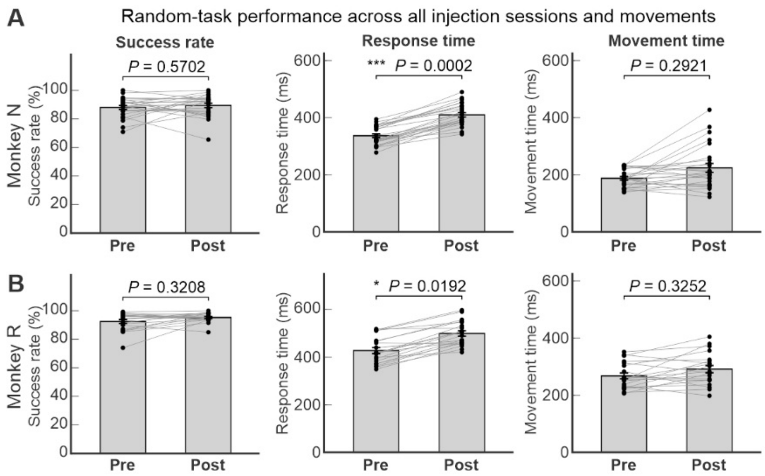


**Fig. S10. Random-task performance across all anisomycin-injection sessions and movement components in each monkey.**

Random-task performance before (Pre) and after (Post) anisomycin injection was analyzed using all available movement components from all injection sessions, without selection based on effect magnitude. Data are shown separately for Monkey N (**A**) and Monkey R (**B**). Columns show percent-correct responses, response time (RT), and movement time (MT). Each point represents one movement/target group from one injection session, and lines connect the corresponding Pre and Post observations. Bars indicate mean ± SE. Predictive responses are not defined for the visually guided Random task. Statistical comparisons were performed using the population analyses described in Methods; exact *P* values are shown above each panel. Percent-correct performance and MT were unchanged in both monkeys, whereas RT increased significantly after anisomycin injection.

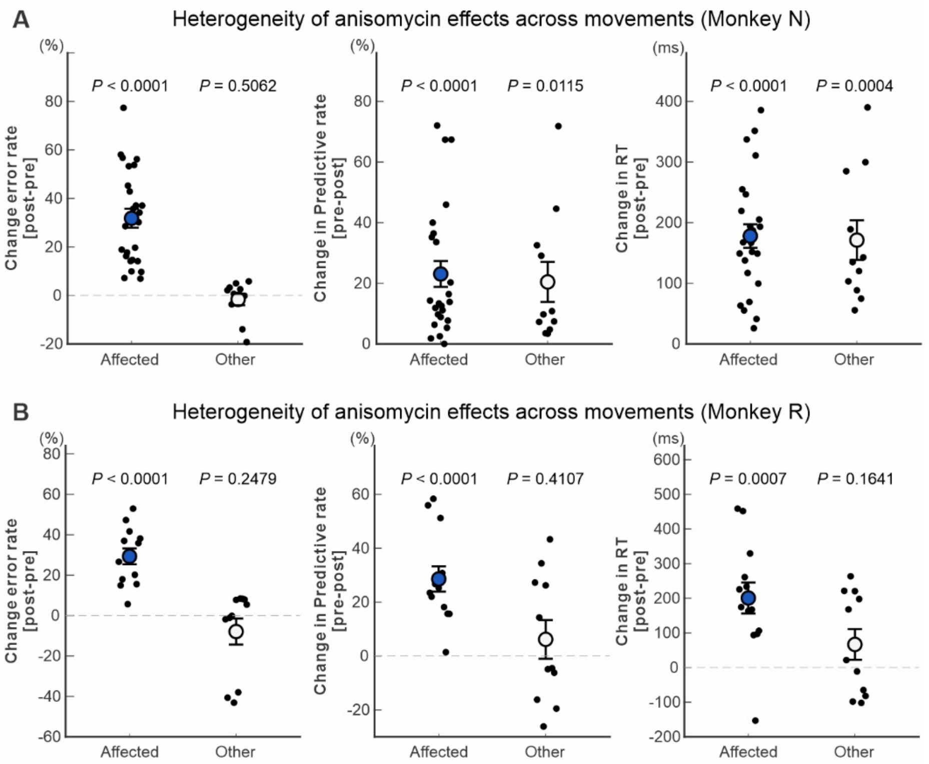


**Fig. S11. Heterogeneity of anisomycin-induced behavioral effects across component movements of the Repeating task.**

Changes in performance from pre- to post-anisomycin injection across all injection sessions are shown separately for Monkey N (**A**) and Monkey R (**B**). Movement components showing a significant pre–post increase in error rate were classified as Affected, whereas the remaining component movements were classified as Other. Each black dot represents one movement component from one injection session. Panels show the change in error rate (post − pre), predictive-response rate (pre − post), and response time (RT; post − pre), such that positive values indicate impairment. Large symbols and error bars indicate mean ± SE; blue filled symbols denote Affected movements and open symbols denote Other movements. Dashed horizontal lines indicate no change from pre-injection performance. Statistical values above each group test whether the change differed from zero. Because classification as Affected was based on a statistically significant increase in error rate, the error-rate panels are descriptive of the classification and do not constitute an independent test of movement heterogeneity. RT reflects response selection and initiation rather than movement execution itself. These analyses illustrate the heterogeneity in the magnitude of anisomycin-induced behavioral changes across component movements.

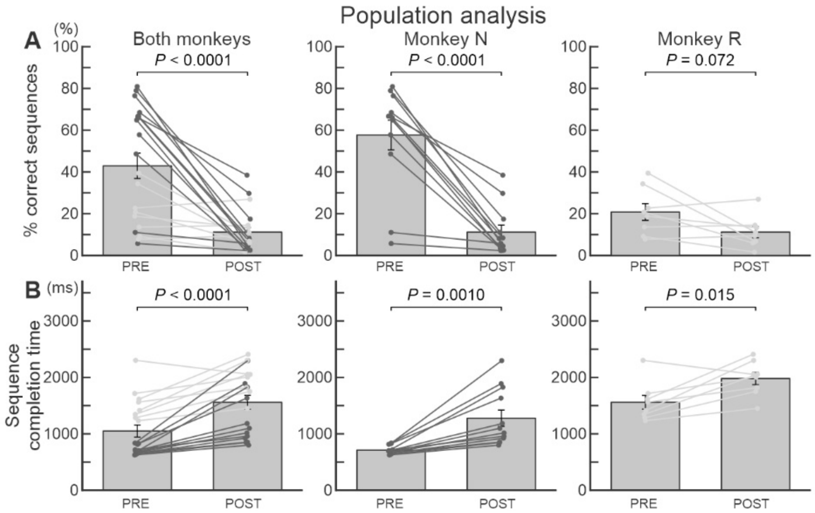


**Fig. S12. Population-level effects of anisomycin on complete-sequence performance.** Complete three-movement sequences were analyzed across all anisomycin-injection sessions. A sequence was scored as correct only when all three component movements were completed correctly on the first attempt.

**(A)** Percentage of correctly completed sequences before (PRE) and after (POST) anisomycin injection. Anisomycin reduced complete-sequence accuracy in the combined dataset and in Monkey N (paired *t* test, $P < 0.0001$ for both). In Monkey R, the paired comparison across sequence-specific observations did not reach significance ($P = 0.072$). However, a complementary trial-level binomial mixed-effects analysis confirmed a significant reduction in the odds of correct sequence completion in Monkey R (21.1% PRE vs. 13.5% POST; odds ratio = 0.58, 95% CI = 0.49–0.69).

**(B)** Sequence completion time for correctly completed sequences before and after anisomycin injection. Completion time increased significantly in the combined dataset ($P < 0.0001$), Monkey N ($P = 0.0010$), and Monkey R ($P = 0.015$; paired *t* tests).

Each connected pair represents one sequence from one injection session; bars show mean ± SE. Data are shown for both monkeys combined (left), Monkey N (middle), and Monkey R (right). Together, these analyses show that anisomycin impaired complete-sequence performance across the population, reducing sequence accuracy and prolonging sequence completion time.

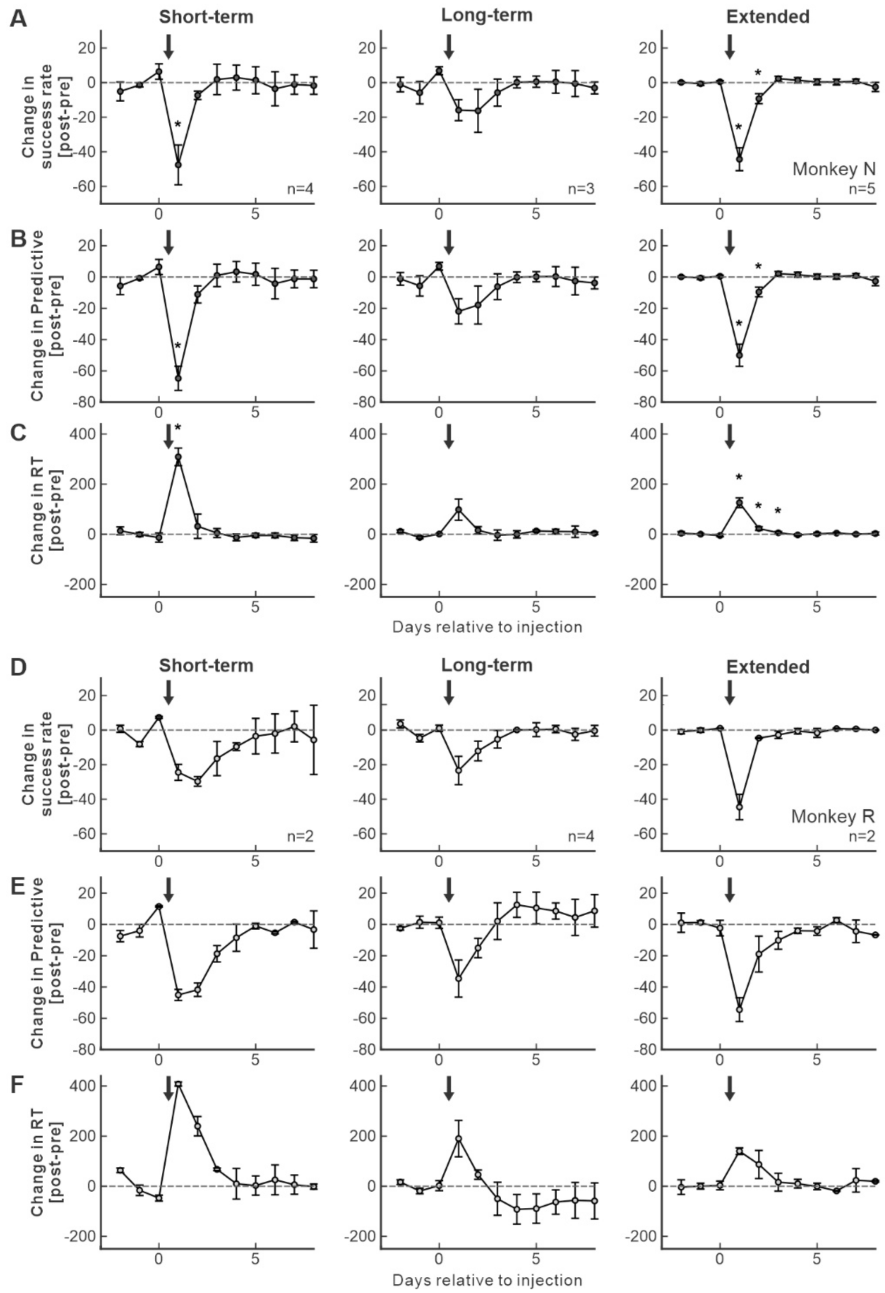


**Fig. S13. Recovery of anisomycin-induced behavioral impairments across training stages in each monkey.** Time courses of changes in Repeating-task performance are shown separately for Monkey N **(A–C)** and Monkey R **(D–F)** for sequences with short-term, long-term, and extended training histories. Values were normalized to the pre-injection baseline and are expressed as post-minus-pre changes in success rate **(A, D)**, predictive-response rate **(B, E)**, and response time (RT) **(C, F)**. Day 0 corresponds to the last pre-injection session; arrows indicate anisomycin injection. Symbols and error bars indicate mean ± SE across injection sessions within each training-stage group. Negative values for success rate and predictive-response rate and positive values for RT indicate impairment. Dashed horizontal lines indicate no change from baseline. Asterisks indicate significant differences from the pre-injection performance ($P < 0.05$; statistical tests as described in Methods). Statistical tests were not performed for conditions represented by fewer than three injection sessions (Monkey R, short-term and extended training).

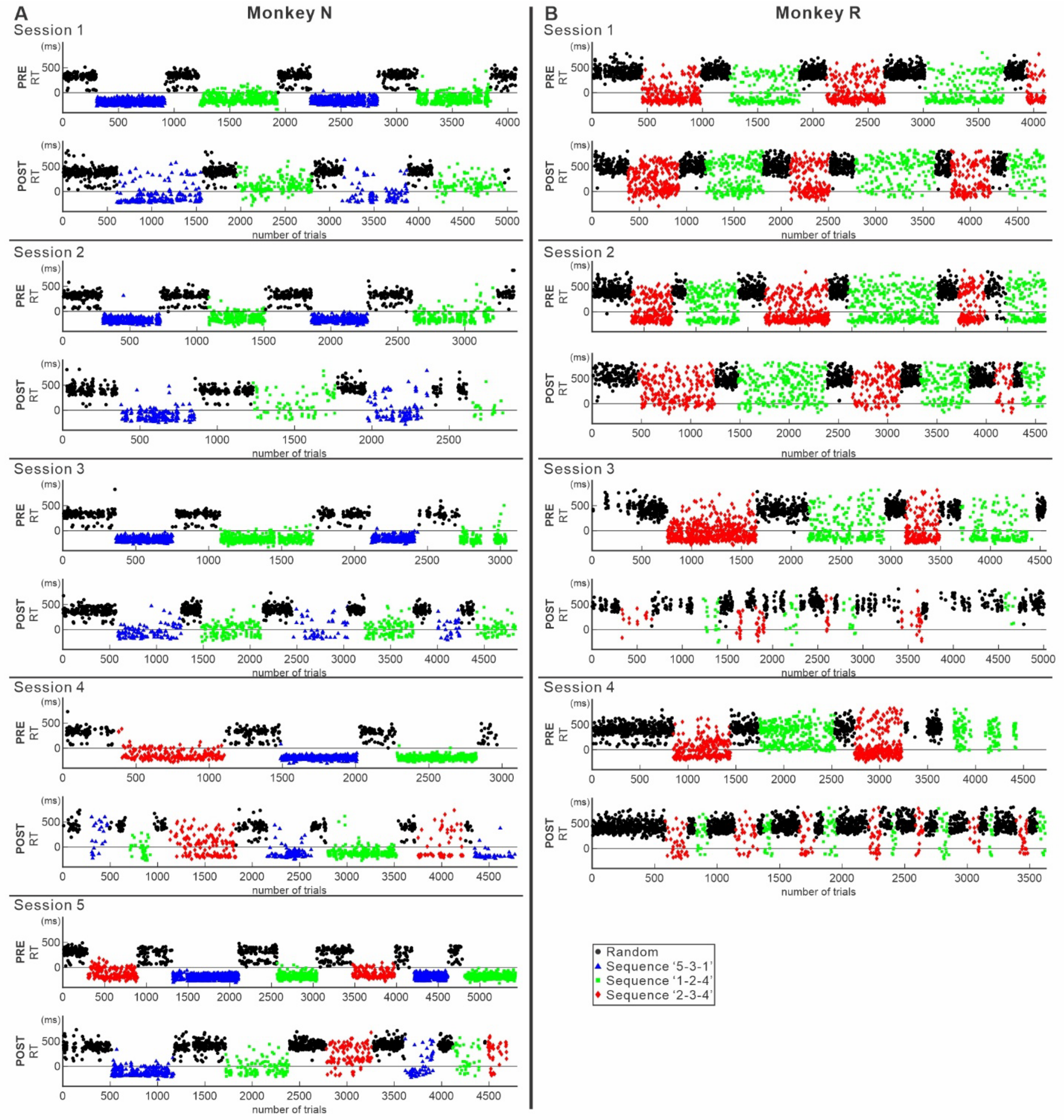


**Fig. S14. Trial-by-trial response times before and after anisomycin injection across all injection sessions in Monkeys N and R.**
**(A)** Monkey N, anisomycin-injection sessions 1–5.
**(B)** Monkey R, anisomycin-injection sessions 1–4.
For each session, response times (RTs) of correct responses are plotted in trial order before injection (PRE, upper plot) and 20–24 h after injection (POST, lower plot). Black symbols indicate the Random task. Repeating-task sequences are shown as sequence 5-3-1 (blue triangles), sequence 1-2-4 (green squares), and sequence 2-3-4 (red diamonds). Each symbol represents one correct movement response; the x-axis shows trial number within the daily session. The monkeys performed Random and Repeating tasks in alternating blocks, resulting in the blockwise organization visible across each session. RTs in the Repeating task were

frequently negative before injection because movements could be initiated predictively before presentation of the visual instruction cue. After anisomycin injection, RT distributions shifted toward longer values in many Repeating-task blocks, although the magnitude and temporal evolution of the effect varied across sequences, movements, and sessions.

In Monkey R, neural activity was recorded after the injection during sessions 3 and 4. Task blocks were switched more frequently in these sessions to optimize sampling across behavioral conditions during neural recording.

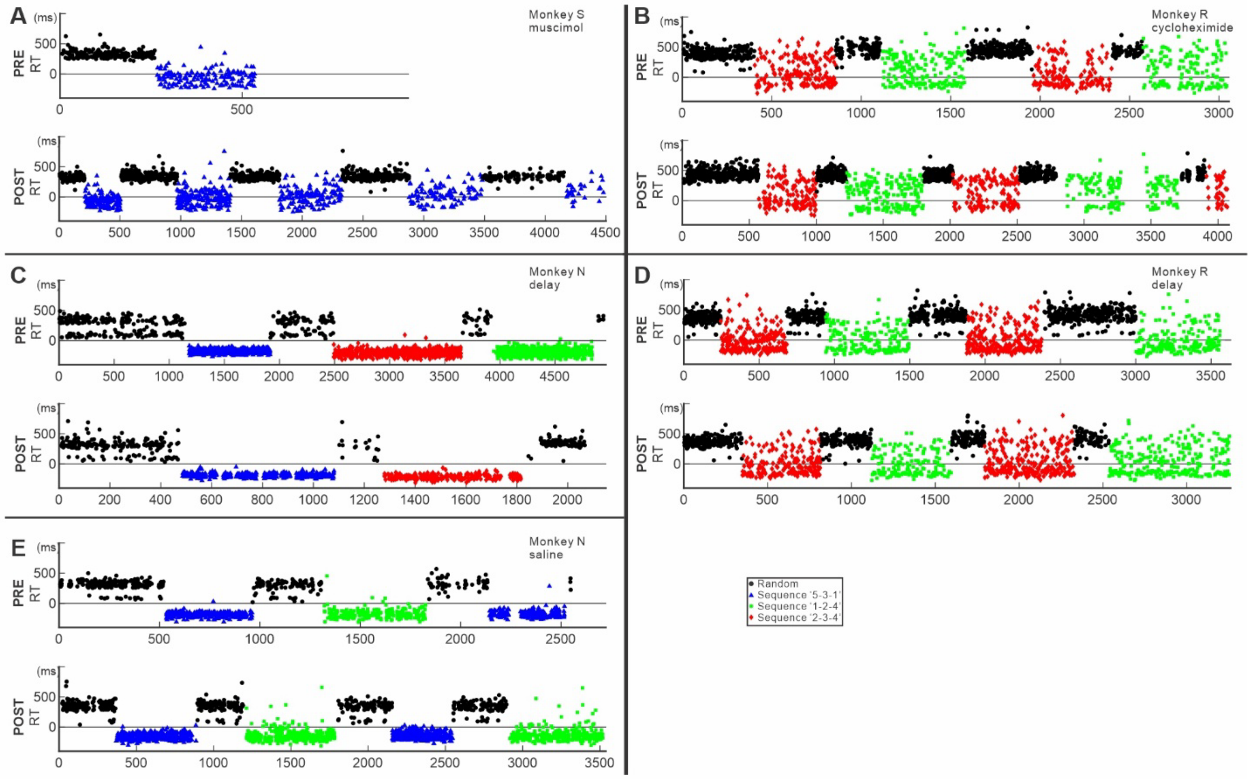


**Fig. S15. Trial-by-trial response times in control injection and delayed-testing sessions.** Response times (RTs) of correct responses during the Random task (black circles) and Repeating tasks (sequence 5-3-1, blue triangles; sequence 1-2-4, green squares; sequence 2-3-4, red diamonds) are plotted in trial order before injection (PRE, upper plots) and during the corresponding post-injection test session (POST, lower plots). Each symbol represents one correct movement response.

**(A)** Muscimol injection control in Monkey S. Muscimol effects were movement dependent and were strongest for movements that showed impaired reaching in both the Random and Repeating tasks. Because many post-injection trials for these movements were errors, and the present plots include only correct-response RTs, the most strongly affected movements are underrepresented in the POST RT plot.

**(B)** Cycloheximide injection in Monkey R. Cycloheximide inhibits protein synthesis through a mechanism distinct from anisomycin.

**(C)** Anisomycin injection in Monkey N followed by delayed testing. The monkey did not perform the task during the post-injection interval and was tested five days after injection.

**(D)** Anisomycin injection in Monkey R followed by delayed testing. The monkey did not perform the task during the post-injection interval and was tested three days after injection.

**(E)** Saline injection control in Monkey N.

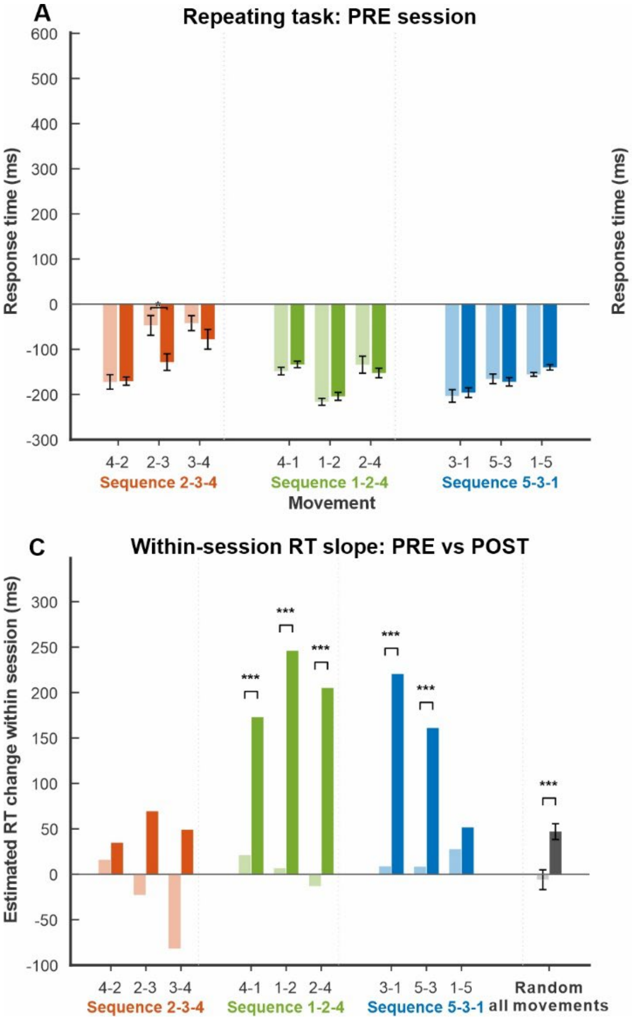


**Fig. S16. Within-session evolution of response time before and after anisomycin injection and delayed testing.**

**(A)** PRE-injection Repeating-task session from representative anisomycin session A5 (Monkey N). For each component movement in sequences 2-3-4, 1-2-4, and 5-3-1, bars show mean ± SE RT for the first 20 (light color) and last 20 (dark color) correct first-attempt responses. First-versus-last comparisons were performed with two-sided Welch two-sample *t* tests, with Holm correction across the three movements within each sequence.

**(B)** POST-injection Repeating-task session from the same experiment, analyzed in the same manner as in A.

**(C)** Comparison of the change in RT over the course of the session between PRE and POST sessions (PRE, light color; POST, dark color). For each movement, a single within-session RT slope was estimated to summarize how RT changed from the beginning to the end of that session. To calculate this slope, correct first-attempt responses were ordered chronologically and their positions within the session were normalized from 0 (first response) to 1 (last response), and RT was regressed on this normalized response order. PRE–POST differences in slope were tested using a linear model containing normalized response order, condition (PRE or POST), and their interaction. The interaction term tested whether the POST slope differed from the PRE slope, with movement-wise *P* values Holm-corrected across the three movements within each

sequence. An overall Repeating-task analysis used an analogous model with movement included as a categorical fixed effect. For the Random task, all available movements were included in an analogous model with movement as a categorical fixed effect; bars at right show mean ± SE movement-wise slopes. Positive slopes indicate increasing RT over the session, whereas negative slopes indicate decreasing RT. Across the complete nine-session anisomycin dataset, the PRE–POST slope interaction was significant at the population level (trial-level linear mixed-effects model: $\beta = 46.8$ ms, 95% CI [28.5, 65.0], $P < 0.0001$; complementary paired comparison of session-level pooled slopes, $P = 0.032$).
**(D)** Delayed-test control session D in Monkey N. Post-injection task performance was assessed five days after anisomycin injection (Fig. S15C). Error rate (top) and RT (bottom) are shown for Repeating and Random trials before and after the delayed test. Bars show mean ± SE. No significant PRE–POST deterioration was detected in either task. Only correct first-attempt responses with valid RT values were included in the RT analyses.

**Movie S1**. Performance of the Random and Repeating sequence ‘2-3-4’ before anisomycin injection. Data from session A5 of Monkey N. The red asterisk indicates the target to be touched, whereas the yellow circle indicates the target touched by the monkey.

**Movie S2**. Performance of the Random and Repeating sequence ‘2-3-4’ after anisomycin injection. Data from session A5 of Monkey N. The red asterisk indicates the target to be touched, whereas the yellow circle indicates the target touched by the monkey.

**Movie S3**. Performance of the Repeating sequence ‘2-3-4’ after anisomycin injection shown in slow motion. Data from session A5 of Monkey N. The red asterisk indicates the target to be touched, whereas the yellow circle indicates the target touched by the monkey.
